# Automatic detection of complex structural genome variation across world populations

**DOI:** 10.1101/200170

**Authors:** Bo Zhou, Joseph G. Arthur, Hanmin Guo, Christopher R. Hughes, Taeyoung Kim, Yiling Huang, Reenal Pattni, HoJoon Lee, Hanlee P. Ji, Giltae Song, Dean Palejev, Xiang Zhu, Wing H. Wong, Alexander E. Urban

## Abstract

Complex structural variants (cxSVs), e.g. inversions with flanking deletions or interspersed inverted duplications, are part of human genetic diversity but their characteristics are not well delineated. Because their structures are difficult to resolve, cxSVs have been largely excluded from genome analysis and population-scale association studies. To permit large-scale detection of cxSVs from paired-end whole-genome sequencing, we developed Automated Reconstruction of Complex Variants (ARC-SV) using a novel probabilistic algorithm and a machine learning approach that leverages the new Human Pangenome Reference Consortium diploid assemblies. Using ARC-SV, we resolved, across 4,262 human genomes spanning all continental super-populations, 8,493 cxSVs belonging to 12 subclasses. Some cxSVs with population-specific signatures are shared with Neanderthals. Overall cxSVs are significantly enriched in regions prone to recombination and germline *de novo* mutations. Many cxSVs mark phenotypic hotspots (each significantly associated with ≥ 20 traits) identified in genome-wide association studies (GWAS), and 46.4% of all significant GWAS-SNPs catalogued to date reside within ±125 kb of at least one cxSV locus. Common SNPs near cxSVs show significant trait heritability enrichment. Genomic regions affected by cxSVs are enriched for bivalent chromatin states. Rare cxSVs are enriched in neural genes and loci undergoing rapid or accelerated evolution and recently evolved *cis*-regulatory regions for human corticogenesis. We also identified 41 fixed loci where divergence from our most recent common ancestor is via localized cxSV. Our method and analysis framework allow for the accurate, efficient, and automatic identification of cxSVs for future population-scale studies of human disease and genome biology.

## INTRODUCTION

Structural variation (SV) analysis though whole-genome sequencing (WGS) is essential to the study of genome biology and disease etiology (Lappalainen et al. 2019). SVs rearrange genome sequences ≥ 50 bp and, though individually rarer than single-nucleotide variants (SNVs) and indels (The 1000 Genomes Project Consortium et al. 2012), account for a much greater portion of the DNA sequence variation between individuals (Pang et al. 2010). Due to their wide size spectrum and complexity, SVs can affect genome function through numerous mechanisms (Stankiewicz and Lupski 2010) and inordinately (by effect size) impact heritable gene expression and disease phenotypes (Stranger et al. 2007; Weischenfeldt et al. 2013; Chiang et al. 2017).

While SV detection methods based on long-read sequencing technologies are rapidly developing (Logsdon et al. 2020), large cohort and population-scale studies (Wonkam 2021; Gaziano et al. 2016; Saunders et al. 2019) continue to rely on paired-end short-read sequencing as it remains the most mature, well-supported, standardized, and cost-effective way to gather genome information (Abel et al. 2020; Collins et al. 2020). To date, standard genome SV analysis focuses almost exclusively on simple types of SVs identified by a single breakpoint, *i*.*e*., by a pair of genomic locations adjacent in the sample but not the reference, such as deletions, tandem duplications, inversions, and/or translocations (Guan and Sung 2016; Kosugi et al. 2019). Detection of these simple-SV breakpoints is theoretically straightforward but difficult in practice (Escaramís et al. 2015; Cameron et al. 2019), and investigators typically apply multiple methods together with heuristic filters (Sudmant et al. 2015; Parikh et al. 2016; Mohiyuddin et al. 2015; Chaisson et al. 2019; Zhou et al. 2018b). However, SVs with complexity beyond the scope of most detection algorithms are abundant in human genomes and have been observed in a variety of phenotypic contexts including healthy, somatic, and cancer tissue (Quinlan and Hall 2012; Sudmant et al. 2015; Collins et al. 2017; Sekar et al. 2020; Collins et al. 2020; Byrska-Bishop et al. 2022; Fujimoto et al. 2021).

Our definition of a complex SV (cxSV) is any genome rearrangement that involves multiple breakpoints not reducible to non-overlapping deletions, tandem duplications, novel insertions, or inversions. More specifically, these cxSVs are localized, with all rearranged reference segments originating from a small to medium-sized (< 2 Mb) genomic region (Table S1, Figure 1, Supplementary Figure S1), unlike chromosomal-scale rearrangements or chromothripsis (Stephens et al. 2011; Forment et al. 2012). Although, complex SVs have been shown to cause Mendelian disorders and associate with other genomic disorders (Sanchis-Juan et al. 2018; Carvalho and Lupski 2016), their roles in genome biology, disease, and evolution are still very poorly understood. This is primarily due to the technological challenges in automatically resolving their sequence structures from standard WGS data where cxSVs are either inadequately represented by a single SV or incorrectly detected as multiple overlapping SVs (Sudmant et al. 2015) (Supplementary Figure S2). While cxSVs have been studied by focused breakpoint analysis with semi-manual inspection of paired-end mapping patterns (Yalcin et al. 2012; Sekar et al. 2020; Collins et al. 2020) and long sequencing reads and mate-pairs (Sudmant et al. 2015; Collins et al. 2017; Frith and Khan 2018; Cretu Stancu et al. 2017; Sedlazeck et al. 2018; Zhou et al. 2019; Lin et al. 2022), computationally efficient and accurate methods for their automatic detection and assembly from standard WGS data that are especially suitable for large-cohort or population-scale datasets have been challenging to develop (Collins et al. 2020; Abel et al. 2020; Byrska-Bishop et al. 2022)

**Figure 1.**
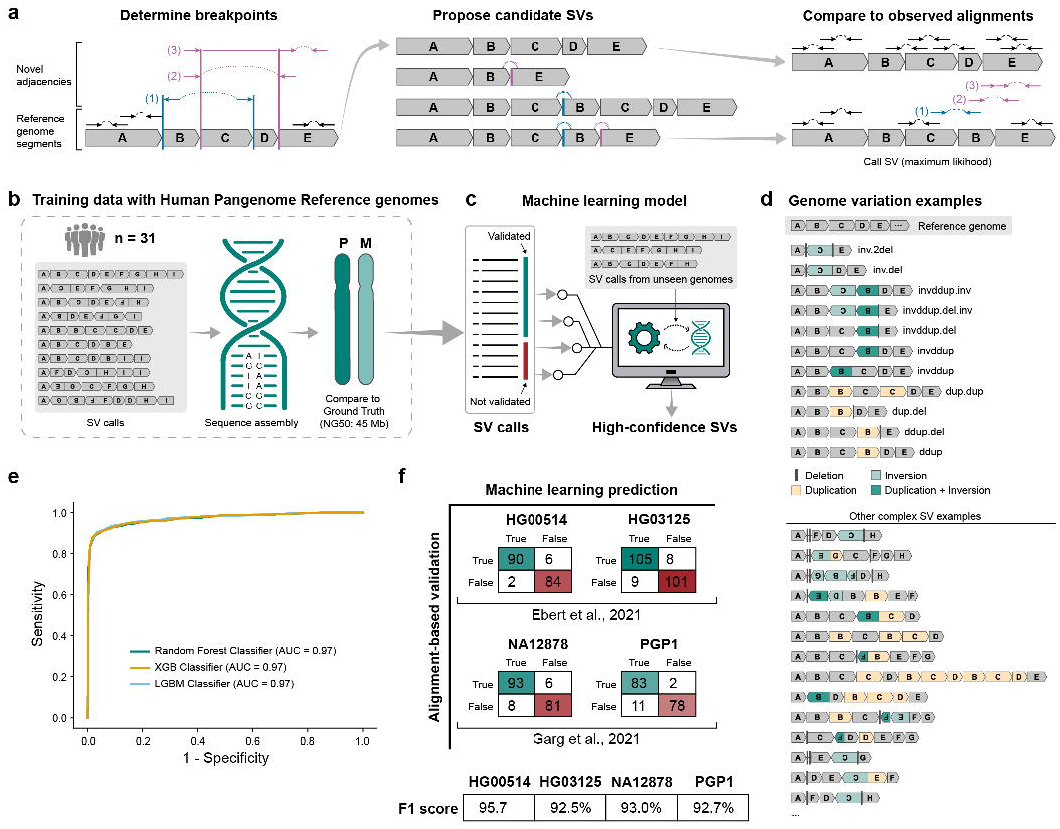
High-level overview of the ARC–SV method. (a) Given a set of paired-end read alignments, candidate breakpoints are determined from discordant read pairs (1 and 2) as well as split reads (3) and soft-clipped alignments. In this case, read 1 suggests the AB and CD breakpoints (orange) while reads 2 and 3 support the BC and CD breakpoints (purple). Possible novel adjacencies (dotted) are then determined e.g. Read 1 suggests an adjacency CB (orange dotted line) while reads 2 and 3 suggest the adjacency BE. Candidate haplotype structures are proposed by enumerating paths through the novel adjacencies. Diploid genotypes are scored using a likelihood model for observed alignments. In this case, the rearrangement ABCBE on one haplotype explains the discordant pairs (1 and 2) and split read (3) that did not match the reference. (b) ARC-SV calls from 31 HPRC individuals were sequence assembled and aligned against their respective HPRC diploid assemblies (Ground Truth) and (c) labeled as “validated” or “not validated”. Features from these calls were then used to train a machine learning model to identify high-confidence SVs from other “unseen” genomes. (d) Examples of complex SVs that can be called by ARC-SV. (e) ROC curve of machine learning models trained to predict cxSV validation in 11 test genomes. (f) Confusion matrices for performance of XGB machine learning model in non-HPRC genomes to predict cxSV validation.

Here, we have developed the novel method, Automated Reconstruction of Complex SVs (ARC-SV), which uses a generative statistical model and a machine learning framework based on diploid genome assemblies (Liao et al. 2022) to accurately detect and automatically sequence assemble an unprecedented spectrum of localized complex structural rearrangements from standard short-read, paired-end WGS data (Figure 1, Table S1, Supplementary Methods). This data-driven, generative approach eliminates any *a priori* assumptions of the types of complex SV structures to be observed in the sample genome. Our method efficiently integrates the automatic detection and assembly of cxSVs to standard WGS analysis. To catalogue and to decipher the characteristics of complex structural variation in natural human genetic diversity, we then applied ARC-SV to 4,262 human genomes that span across all 8 continental super-populations across the world (Table 1). We then investigated the characteristics of their distribution across the genome, their proximity to GWAS signatures, the chromatin states of affected regions across various tissues, their enrichment in genes and regulatory regions, and their occurrence in genome regions under rapid or accelerated evolution including those that diverged from our closest primate relative via cxSV, as well as cxSVs shared between modern and archaic humans.

## RESULTS

### ARC-SV detection of cxSVs using a probabilistic and machine learning framework

At a given stretch of the genome, there are in principle numerous possible cxSV structures (Quinlan and Hall 2012), each producing its own unique signatures of read depth, mapped read orientations, insert sizes, and split alignments (Sekar et al. 2020). The observed signature will depend on sequencing library characteristics (especially the insert size distribution) in addition to the structure and size of the SV. Several previous methods (Collins et al. 2017, 2020; Byrska-Bishop et al. 2022) attempt to classify cxSV structures by direct comparison to these mapping signatures. Here, to achieve more general detection of cxSVs, ARC-SV simultaneously detects and genotypes SVs by integrating all reads into a probabilistic model (including reads that match the reference) which can be used to score proposed candidate diploid rearrangements (Figure 1a) in contrast to other SV analysis methods that only interrogate breakpoint-supporting reads (Collins et al. 2020; Byrska-Bishop et al. 2022) (see Supplementary Methods for detailed description of probabilistic model). Briefly, given a standard paired-end read alignment dataset, candidate breakpoints are first determined from soft-clipped reads, split-read alignments, and clustering of discordant read pairs (determined from library insert-size distribution and coverage that are unique to each sequencing library) (Supplementary Figure S3). Breakpoint intervals determined from discordant read pair clusters are refined based on a probabilistic model and merged with those determined from soft-clipped/split reads. Based on breakpoint locations, ARC-SV then defines blocks of genome segments that are rearranged in the sample genome and constructs adjacency graphs that represent reference or novel (non-reference) adjacencies between genomic segments. Candidate haplotype structures are then proposed by enumerating paths through the adjacency graph and evaluated based on paired-end genome alignment likelihood modeling (Figure 1a, Supplementary Methods). To make complex SVs structures more interpretable, we devised our own convention of describing all cxSVs using an alphabetical notation (e.g. ABCDE → ABC’B’E where B segment is duplicated in an inverted manner between C and E, C segment is inverted, and D segment is deleted (Figure 1, Table S1). In this process of detecting cxSVs, ARC-SV also automatically detects, genotypes, and sequence assembles canonical simple SVs, e.g. deletions, inversions, tandem duplications (Table S1).

To further increase the precision and accuracy of the SV calls, i.e. to identify high-confidence calls from the ARC-SV probabilistic model, we integrated a machine learning framework trained to distinguish SV calls that validate against established “ground truths” (Figure 1b-c). Here, for the “ground truth” set, we leverage 42 new state-of-the-art diploid assemblies of diverse individuals released by the Human Pangenome Reference Consortium (HPRC) (Liao et al. 2022) (Table S2). For training and testing of our machine learning models, the 42 diploid assemblies are split into training (n=31) and test (n=11) sets stratified by genome ancestry (Table S2).

For all SVs, and perhaps especially for cxSVs, it is essential that calls are validated independently. Validation of simple SV calls is traditionally conducted by comparison to an existing gold set of predictions (Zook et al. 2020) where a SV prediction is typically said to be validated if it has high overlap to one of the “true” SVs. The commonly-used criterion is that the overlap between the predicted and true SVs is at least 50% of the larger one’s size — this is referred to as 50% reciprocal overlap (Rausch et al. 2012; Hormozdiari et al. 2009; Zhou et al. 2018b). Applying the same kind of methodology to cxSV predictions is infeasible since gold-standard cxSV call sets are not available. Here, we validate SV calls from an individual by direct sequence match to the individual’s diploid genome assembly (“ground truth”) via first sequence assembly of detected SVs followed by sequence alignment (Figure 1b-c).

ARC-SV automatically constructs a sequence assembly for each SV call with 1kb of flanking sequences on each side. For genomes (n=42) with HPRC assemblies (Table S2), SVs called from standard paired-end WGS of the same genomes are sequence assembled and aligned to the HPRC genome assemblies. Before alignment, predicted SV sequences are modified such that only 150 bp of flanking sequences are included. This is so that flanking sequences serve only the role of anchor and have minimal effect on the overall alignment score of the prediction sequence. Using an established validation criteria of cxSV detection using long-reads (Sedlazeck et al. 2018), a cxSV call is validated if its sequence prediction of rearranged genome segments matches the “ground truth” via complete and continuous sequence alignment (Supplementary Methods). See Figure 2 for examples. Compared to commonly used validation methods such as *in silico* simulation and overlap based on SV call set and/or alignment spans, our validation method, although significantly more stringent, is better suited to train accurate machine learning models. ARC-SV cxSV calls from each HPRC genome (Table S2) based on ∼30×-coverage paired-end WGS (Byrska-Bishop et al. 2022) were validated against the diploid genome assemblies and labeled based on validation outcome (Table S3). Of the initial ARC-SV cxSV calls prior to machine learning, 54% (n=4,221) of total cxSV calls across the HPRC genomes were validated and 46% (n=3,665) were not (Table S3).

**Figure 2.**
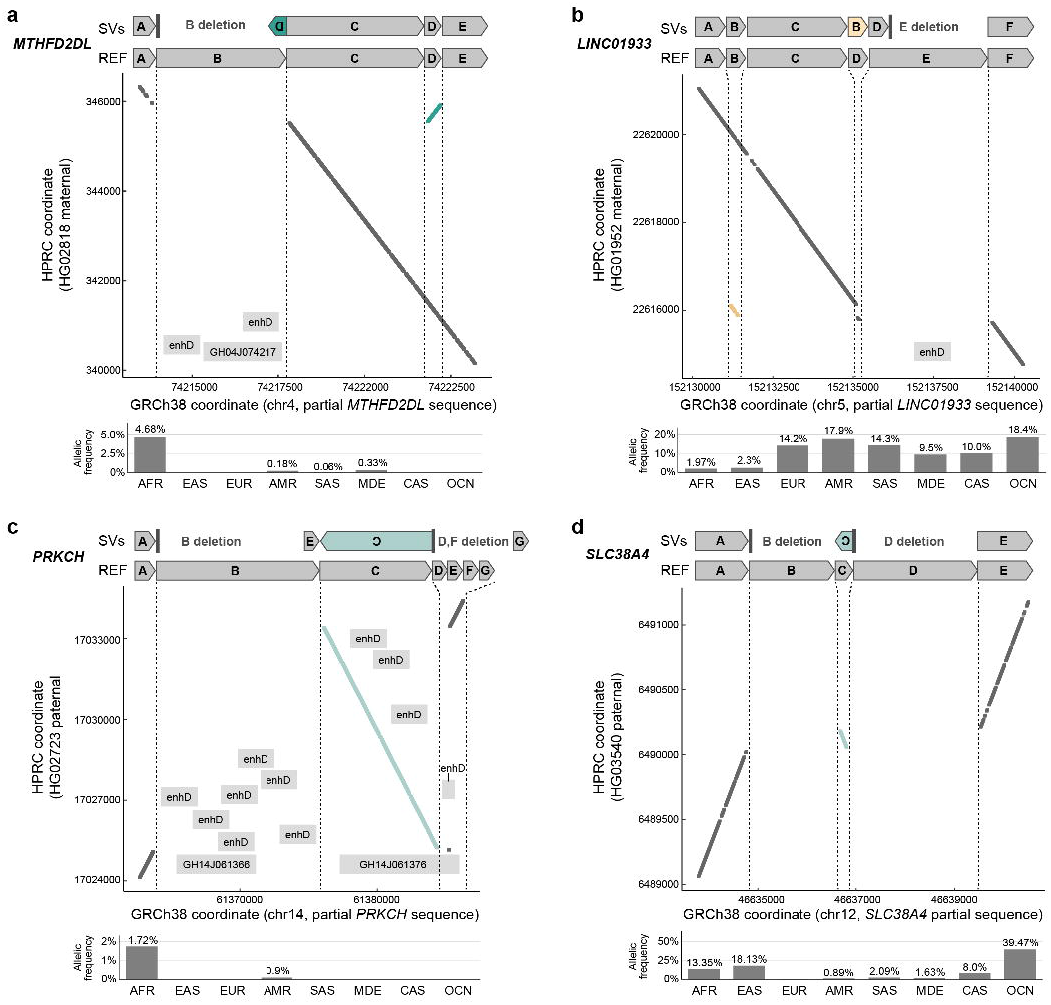
Examples of cxSVs validated in HPRC assemblies and with population-specific signatures. (a-d) Plots mapped genomic locations of unique k-mers (31bp) in GRCh38 (x-axis) and respective HPRC assemblies (y-axis) at loci with cxSV identification by ARC-SV. ENCCODE cCREs with distal enhancer-like signatures indicated by “enhD”. GH#s represent GeneHancer annotations. Bar plots represent AF (%) in each superpopulations.

Conventionally, high-confidence SV calls are typically filtered based on metrics such as the number of supporting reads or the depth of coverage in surrounding regions (Layer et al. 2014; Rausch et al. 2012; Chen et al. 2016). While this approach is interpretable and easy to communicate, it does not make full use of the data and requires the user to manually specify thresholds, often without any recommendations. Our alignment-based validation approach (Sedlazeck et al. 2018) provides the opportunity for a more careful study of the accuracy of SV prediction.

Our machine learning model predicts SV validation given 48 features (Table S4) transformed from the default output table of the ARC-SV probabilistic model (Table S3, Supplementary Methods). From these transformed features, we then trained random forest, extreme gradient boosting, and light gradient boosting models (Figure 1b-e, Source Data). Maximizing for precision, we trained a separate model for each SV class (cxSVs, deletions, tandem duplication, and inversion). The optimal machine learning model for predicting high-confidence cxSVs is an extreme gradient boosting (XGBoost) model achieving 95.7% precision and 91.4% recall on the test set (Table S5, Figure 1e). Machine learning models for simple SVs were also trained using the same alignment-based validation approach against HPRC diploid assemblies: deletions (XGBoost, 99% precision, 95.8% recall), tandem duplications (XGBoost, 97.8% precision, 90.9% recall), and inversions (random forest, 96.2% precision, 96.2% recall) (Tables S4-5).

### Additional validation of ARC-SV detection of cxSVs

To further validate our method, we also performed alignment-based validation of high-confidence cxSV calls against other recently published diploid genome assemblies as “ground truths”; these were constructed using different methods from the HPRC assemblies (Liao et al. 2022). We validated our high-confidence cxSV calls (identified via machine learning model) for NA12878 and PGP1 (both of European ancestry) against their respective diploid genomes assembled using DipAsm (Garg et al. 2021) as well as those for HG00514 (East Asian ancestry) and HG03125 (African ancestry) against genomes assembled using a combination of PacBio HiFi and Strand-Seq (Ebert et al. 2021). We observed ≥ 92.5% F1 score across all four genome validations (Table S6, Figure 1f).

Furthermore, we also validated our method for cxSV discovery in somatic tissues. It was demonstrated previously that mosaic complex SVs can arise in the developing human brain likely during cell proliferation in mid-neurogenesis (Sekar et al. 2020). Clonal populations (n=41) were obtained from single neural progenitor cells from the cerebral cortex (FR), parietal cortex (PA) and basal ganglia (BG) regions of three phenotypically normal postmortem human fetuses (Bae et al. 2018); each clone was sequenced via standard WGS to 30× coverage, and through careful manual breakpoint analysis and local assembly, 3 localized intragenic cxSVs (2 of which are within neurodevelopmental genes *SPAST* and *NAV3*) and one inter-chromosomal cxSV were experimentally validated to be mosaic, present in 4 different clones belonging to 4 different brain regions (Sekar et al. 2020). We applied ARC-SV to all 41 clones for mosaic cxSV discovery. We then compared raw cxSV calls across different clones to identify those that may be somatic or those that occur only in subsets of clones within the same tissue of origin and obtained 42 candidates for putative mosaic cxSVs (Table S7) where we then applied our trained machine learning model to filter for high-confidence cxSVs. Our model retained only the 3 localized intragenic cxSVs and in the exact same clones that were experimentally validated (Sekar et al. 2020).

In addition, we also used PCR followed by Sanger sequencing to experimentally validate cxSV calls (n=23) from two pilot genomes that were originally used for methods development: HuRef (Zhou et al. 2018a) and HepG2 (Zhou et al. 2019) (Table S8). Primers were designed to amplify the cxSV in its entirety i.e. all breakpoints in a single reaction (Table S9). For cxSVs called as heterozygous, we ensured that reference alleles (without SV) were also amplified to validate genotype. Sanger sequencing from both forward and reverse primers was performed. We observed validation rates of 90% (9/10) for HuRef and 85% (11/13) and for HepG2 (Table S9).

### Comparison against other short or long read-based cxSV detection methods

Recently, two studies on human structural variation involving the Human Structural Variation Consortium and the Genome Aggregate Database Consortium were published that incorporated detection of complex SVs from standard WGS at a population-scale (Collins et al. 2020; Byrska-Bishop et al. 2022). Both studies used the same core method developed in Collins *et al* (Collins et al. 2020) that incorporates an ensemble of different tools for breakpoint analysis to enable cxSV detection. Here, since we analyzed the same 1000 Genomes Project (1000GP) samples (n=3,202) using the same GRCh38 alignment data as Byrska-Bishop et al (Byrska-Bishop et al. 2022), we benchmarked our cxSV calls against those made by Byrska-Bishop *et al* (Byrska-Bishop et al. 2022) for genomes (n=40) with “ground truths” assemblies (Liao et al. 2022) (Table S2). Following our alignment-based validation strategy, we first sequence assembled the cxSV calls from Byrska-Bishop *et al* (Byrska-Bishop et al. 2022) with 150 bp of reference sequence flanking each side (Source Data) and then aligned these sequence of assembled cxSVs directly against their respective assembled genomes (ground truths) for validation (Supplementary Method). Since we used 31 of these genomes (Table S2) for training our machine learning models (Figure 1b-c,e-f, Table S5), we first benchmarked raw or naïve ARC-SV calls (without machine learning) for this set of genomes where we observed 188 ± 17 (mean, standard deviation) cxSV calls per genome with 53% ± 2.8% validation (Table S3), compared to 135 ± 12 cxSV calls per genome with 22% ± 2.6% validation for Byrska-Bishop *et al* (Byrska-Bishop et al. 2022) (Table S10, Figure S4). For the other 9 benchmarking genomes (Table S2), we observed 99 ± 10 cxSV calls per genome with 95% ± 0.7% validation rate for ARC-SV (with machine learning) (Table S3), compared to 131 ± 9 cxSVs calls per genome with 28% ± 5.6% validation for calls from Byrska-Bishop *et al* (Byrska-Bishop et al. 2022) (Table S10, Figure S5). Of the validated cxSV calls from Byrska-Bishop *et al* across these 40 genomes, 81% ± 6.3% were also called by ARC-SV, and of these, the ARC-SV call was not validated in 3 ± 1 cases per genome (Table S10).

In theory, long reads are better suited for resolving cxSVs as they are more likely to span and phase consecutive breakpoints. We thus compared the performance of ARC-SV to that of a novel cxSV detection method based on deep learning of long-read alignments, SVision (Lin et al. 2022). We first checked whether ARC-SV can resolve the 18 (from NA12878) manually curated cxSVs of six unique structures that was used as the benchmark where all 18 were resolved by SVision (Lin et al. 2022). ARC-SV was also able to resolve 17/18 (94%) cxSVs including all internal breakpoints (Table S11). In the Lin *et al* study, 80 high-confidence cxSV calls from HG00733 were made by SVision for which 20 (55%) were selected for experimental validation where 11 validated (Lin et al. 2022). ARC-SV resolved all 11 experimentally validated events (Table S12) and of the remaining 60 SVision calls, ARC-SV resolved 31/60 (52%) in line with the experimental validation rate for Svision (Table S13). We then sequence assembled the remaining 29/60 Svision calls from HG00733 that were not called by ARC-SV and benchmarked them against the HG00733 diploid assembly (Liao et al. 2022) where 1/29 validated (Table S13), further confirming a validation rate of ∼55% for high-confidence Svision calls. In comparison, ARC-SV made 94 high-confidence cxSVs for HG00733 with 94% validation (Table S3).

### Complex SVs across diverse human populations

After having extensively validated ARC-SV to decipher and automatically resolve cxSVs from standard paired-end WGS data, we applied our method to 4,262 healthy human genomes from a total 134 diverse populations across all 8 continental super-populations to investigate the diversity of complex SVs resulting from natural human genetic variation (Table 1, Figure 3a, Table S14). These genomes are compiled from the 1000 Genomes Project (The 1000 Genomes Project Consortium et al. 2010; Byrska-Bishop et al. 2022), the Human Genome Diversity Project (Bergström et al. 2020), and the Simons Genome Diversity Project (Mallick et al. 2016). Across all genomes, we discovered 8,493 unique cxSVs that span up to 482 kb of the genome with the vast majority spanning between 100 bp to 10 kb (Figure 3b, Table S15). To the best of our knowledge, this is the largest and most comprehensive catalogue of human complex SVs to date. Around 75% of the discovered cxSVs can be categorized into 11 distinct subclasses (inversion deletion, duplication-deletion, deletion-flanked inversions, etc); others are too complex to be verbally described and thus categorized as “Other” (Figure 1d, 3b,c, Table S1,S15).

**Figure 3.**
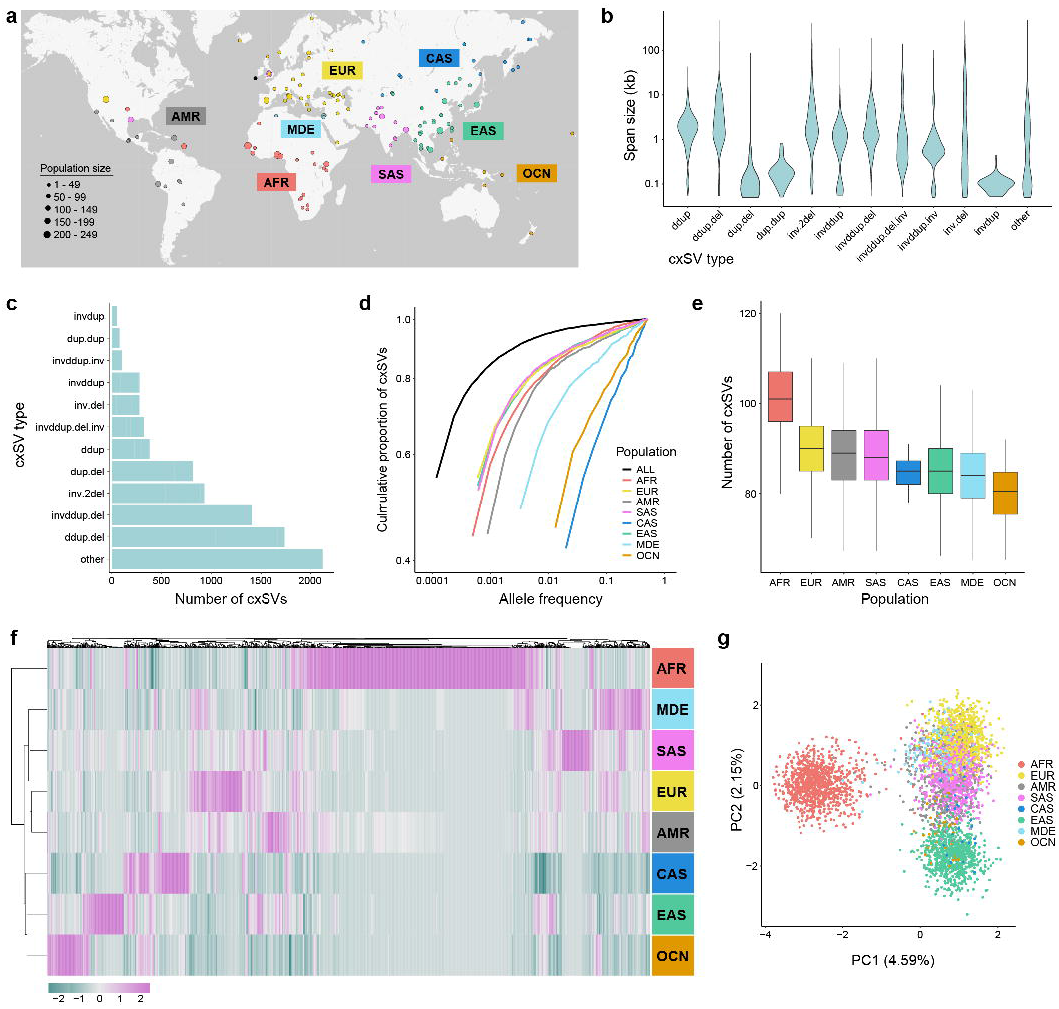
cxSVs across human populations. (a) Geographical distribution and ancestries of 4,262 human genomes studied. (b) Size distribution of 12 classes of cxSVs identified. (c) Total number of cxSVs by class. (d) Cumulative proportion of cxSVs against allele frequencies for different population. (e) Boxplot of number of cxSVs by superpopulation. (f) heatmap showing common cxSV across superpopulations where color scale indicate z-score of allele-frequency. (g). Scatterplot of top two principal components for common cxSVs.

Approximately 16.2% (n=1,377) of these unique cxSVs have an allele frequency ≥ 0.1% and thus considered common, and 7,116 are below 0.1% allelic frequency and considered rare; 4,661 (54.9%) of cxSVs are private to single individuals (Figure 3d, Table S15). Overall, the allele frequencies of cxSVs are lower than that of simple SVs in the human population (one-sided Mann-Whitney test, p-value = 4.2e-14, Figure S6). At an individual level, individuals of AFR descent have on average the highest number (n = 101) of cxSVs compared to other super-populations (Welch’s one-sided two sample t-test, p = 2.9e-305) (Figure 3e). Overall, the majority of unique cxSVs are rare at a population level (Figure 2D), while within any given individual, >90% of the cxSVs will be common events with ∼5-10 private cxSV events with a typical span size of ∼6.2 kb (Table S15). 45.2% of cxSVs overlap protein coding genes, and 66.5% overlapping putative ENCODE regulatory regions (The ENCODE Project Consortium 2012). We also find that 69.4% (956/1377) of common cxSVs (AF ≥ 0.1%) fall within previously determined structural polymorphic genomic regions determined based on the 47 HPRC genomes (Lee et al. 2022).

The majority of common cxSVs show strong signatures of population specificity (Figure 3f). For example, an interspersed inverted duplication with deletion cxSV (AD’CDE) on chromosome 4 (validated in maternal assembly of HG02818, Gambian female, Figure 2a) is common in AFR, AMR, and MDE but rare elsewhere (Table S15). This cxSV resides within *MTHFD2L*, which is associated with cytokine secretion in response to viral infections and vaccines (Kennedy et al. 2012). This cxSV deletes 2 ENCODE candidate *cis*-regulatory elements (cCREs) (E2303693 and E2303696) with distal enhancer-like signatures (The ENCODE Project Consortium 2012) and enhancer (GH04J074217) (Fishilevich et al. 2017) which are bound by key developmental transcription factors such as IKZF1, FOXA1 and FOXA2 (Hammal et al. 2022). Another example is a cxSV (ABCBDF) on chromosome 5 (validated in the maternal assembly of HG01952, Peruvian male, Figure 2b) within *LINC01933* where the allele frequency (AF) is ∼2% in AFR and EAS populations but at 9.5-18% in other populations (Table S15). The deletion of “Block E” within this cxSVs deletes a cCRE (E2422272) with distal enhancer-like signature (The ENCODE Project Consortium 2012) that is bound by ESRRA which interacts with PGC-1 transcription cofactors to regulate the expression of most genes involved in cellular energy production and biogenesis (Schreiber et al. 2004; Hammal et al. 2022). Furthermore, a cxSV (ACE’G) on chromosome 14 (validated in the paternal assembly of HG02723, Gambian female, Figure 2c, Table S15) is found only in AFR (AF=1.7%) and AMR (AF=0.09%) populations and resides within *PRKCH* which encodes a transcription factor predominantly expressed in epithelial tissues and can regulate keratinocyte differentiation (Cabodi et al. 2000). The deletion of Blocks B (12.4kb) deletes 7 cCREs, inversion of Block C affects 3 cCREs, and deletions of Blocks DEF partially deletes cCRE E1719464 (The ENCODE Project Consortium 2012) and enhancer (GH14J061376) (Fishilevich et al. 2017) which are bound by WT1 and IRF4 (Hammal et al. 2022), a key regulatory factor of in the development of human immune cells (Nam and Lim 2016). WT1 exhibit complex tissue-specific and polymorphic imprinting pattern, with biallelic, and monoallelic expression from the maternal and paternal alleles in different tissues (Jinno et al. 1994). Lastly, a cxSV (AC’E) on chromosome 12 (validated in the paternal assembly of HG03540, Figure 2d) within *SLC38A4*, a tumor suppressor gene (Li et al. 2021), is common in all populations except EUR with the highest AFs in OCN, EAS, AFR, and CAS populations.

For common cxSVs in general, principal component analysis (PCA) shows distinct clusters for diverse continental super-populations and that the distance and spatial relationship between the clusters reflect and correlate with geographical distances (Figure 3g). Batch effect between source datasets is minimal (Supplementary Figure S7). Compared to other super-populations, AFR has the most population-specific cxSVs, which is consistent with AFR having the highest degree of genetic diversity (Figure 3e-g, Table S15). Moreover, we investigated the gene associations of cxSV regions with top loadings in the top two principal components that distinguish different populations (Supplementary Figure S8). We find that genes located near the top cxSVs contributing to population-specific signatures are enriched for those linked to the distribution of body fat and blood osmolarity (Supplementary Figure S8).

### Enrichment analysis of cxSVs in recombination initiation and DNM hotspots

In addition to world population, we examined how cxSVs are distributed across the genome (Figure 4a). We observed a tendency for cxSVs to be clustered toward chromosome ends (Figure 4a) where these regions have been associated with elevated local mutation rates (Nesta et al. 2021; Audano et al. 2019) and higher recombination rates (Jensen-Seaman et al. 2004). The formation of structural variation in general has often been attributed to the improper repair of double-strand DNA breaks (Carvalho and Lupski 2016). To investigate whether cxSVs are enriched in recombination hotspots, we performed permutation test and observed singificant overlap between cxSVs and genome recombination initiation double-stranded break hotspots (Pratto et al. 2014) (enrichment = 1.33, p < 1.0e-5) (Figure 4a).

**Figure 4.**
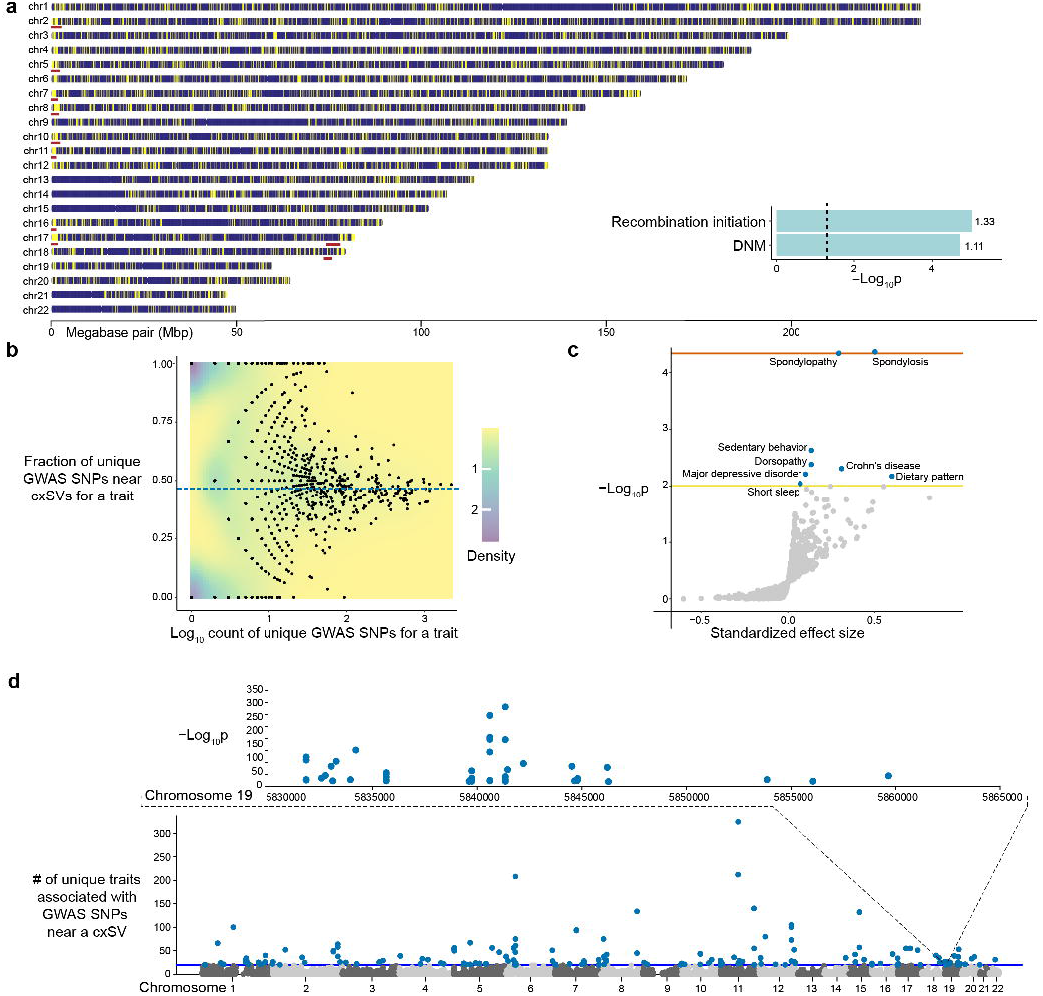
cxSV locations across the genome and their proximity to significant GWAS SNPs. (a) Yellow bars indicate location of cxSV along the chromosome. High density of cxSVs at sub-telomeric regions marked by red. Enrichment p-value and fold indicated by bar plot. (b) Scatter-plot of fraction of unique GWAS SNPs near cxSVs versus count of unique GWAS SNPs for trait; the blue dashed line indicates the fraction across all traits (46.4%). Color scale indicates probability density. (c) heritability enrichment of common SNPs near cxSVs; yellow line indicates nominal p-value=0.01; orange line indicates p-value=0.05 after Bonferroni correction. (d) Manhattan plot of phenotypic hotspots (each is significantly associated with ≥ 20 traits). Example of a phenotypic hotspot chr19:5831855-5859707. Each dot represents a GWAS SNP with the y-axis indicating the -log10 (p-value) from the GWAS.

To test whether cxSVs are enriched in regions with elevated mutation rates, we performed permutation test on the overlap between all detected cxSVs (Table S15) and germline *de novo* mutations (DNMs) from healthy controls annotated in the Gene4Denovo database (Zhao et al. 2020). As DNMs are only present in the offspring and absent in the parents, they occur within a single generation and thus are subjected to minimal selective pressure, their distribution may largely reflect the germline mutation rate (Murat et al. 2022). We observed significant overlap (enrichment = 1.11, p = 2.0e-5) suggesting that cxSVs have a tendency to occur in regions with elevated mutation rates (Figure 4a).

### cxSVs near GWAS SNPs

To gain insight into how cxSVs may affect complex traits and diseases, we integrated cxSVs with GWAS. Most GWAS to date focus only on the associations of common SNPs. To assess the utility of cxSVs in GWAS, we first mapped common SNPs to cxSVs using the gnomAD-SV linkage disequilibrium (LD) map which contains SNVs in strong LD with gnomAD-SVs (R2 ≥ 0.8) (Collins et al. 2020) (Supplementary Figure S9). In the gnomAD-SV LD map, there are 564,676 unique pairs of gnomAD-SVs and SNVs; 506,511 (89.7%) of which have distances less than 125 kb. For AFR individuals, 215,240 (90.5%) pairs are within 125 kb; for EUR individuals, 291,271 (89.1%) pairs are within 125 kb. Informed by theses distributions of LD-distances, we mapped a common SNP in the GWAS Catalog (p-value ≤ 5e-8) (Sollis et al. 2023) to a cxSV if their distance is within 125 kb (Methods).

Among the 40,377 unique SNPs in the GWAS Catalog (henceforth “GWAS SNPs”), 18,732 (46.4%) are mapped to at least one cxSV (Table S16). For the 7,194 unique cxSV intervals (Table S15), 4,121 (57.3%) of which contain at least one mapped GWAS SNP (within ± 125 kb). Among the 2,811 unique GWAS traits, 2,025 (72.0%) have at least one SNP mapped to a cxSV (Figure 4b, Table S16). For these 2,025 traits, 1,572 (77.6%) have higher fractions of GWAS SNPs mapped to cxSVs (Figure 4b) where 29 of show significance after Bonferroni correction (one-sided Binomial p-value ≤ 0.05/2025). Many cxSVs are mapped to GWAS SNPs associated with multiple traits (median: 3; Figure 4d, Table S16). We define a cxSV as a “phenotypic hotspot” if it is mapped to GWAS SNPs associated with ≥ 20 unique traits (Figure 4d), corresponding to the top 5% of trait counts (Table S16). For example the cxSV (AC’E) on chr19:5831855-5859707 is associated with 20 unique GWAS traits (Table S16, Figure 4d) such as cancer biomarkers, heart rate variability, and N-glycan, which is related to brain development (Medina-Cano et al. 2018). The “hottest” phenotypic hotspot contains GWAS SNPs associated with 318 unique traits (Table S16, Figure 4d). The number of unique traits associated with such mapped GWAS SNPs is uncorrelated with the size of cxSV (Pearson R: 0.004; p-value: 0.75). Altogether, our results show that cxSVs are mapped an excess of GWAS SNPs than expected by chance.

In addition to mapping GWAS SNPs to cxSVs, we also assessed if common SNPs (MAF ≥ 0.05) mapped to cxSVs could explain a much larger proportion of complex trait heritability than one would expect by the number of these SNPs (i.e., heritability enrichment (Finucane et al. 2015)) across 1,087 GWAS datasets (Zhu et al. 2022). While significant heritability enrichment (estimate: -0.006; p-value: 0.99) was not observed for meta-analysis estimates across these datasets, we obtained significant heritability enrichments in GWAS of spondylosis (estimate: 0.504; p-value: 4.31e-05) and spondylopathy (estimate: 0.296; p-value: 4.56e-05) that passed the stringent Bonferroni correction (p-value ≤ 0.05/1087), as well as enrichments in GWAS of major depressive disorder, sedentary behavior, dorsopathy, short sleep, dietary pattern, and Crohn’s disease at the nominal significance level (p-value ≤ 0.01; Figure 4c, Supplementary Table S17).

### cxSVs and chromatin states

Since a significantly high fraction of GWAS SNPs are mapped to cxSVs (Figure 4b) than expected by chance and GWAS SNPs are enriched in non-coding regulatory regions (Maurano et al. 2012; Zhu et al. 2021), we then investigated how cxSVs might affect regulatory regions across various tissue types by assessing their enrichment and depletion across 4 regulatory chromatin-state categories and across 127 reference epigenomes (Roadmap Epigenomics Consortium et al. 2015). We find that cxSV regions are enriched for bivalent chromatin regions that encompass bivalent/poised TSS, Flanking Bivalent TSS/Enhancer and Bivalent Enhancer while simultaneously depleted for transcriptionally active chromatin states and enhancers (Figure 5, Table S18). This pattern of enrichment and depletion mirror closely to that of a similar analysis for the fastest evolving regions in the human genome (Mangan et al. 2022).

**Figure 5.**
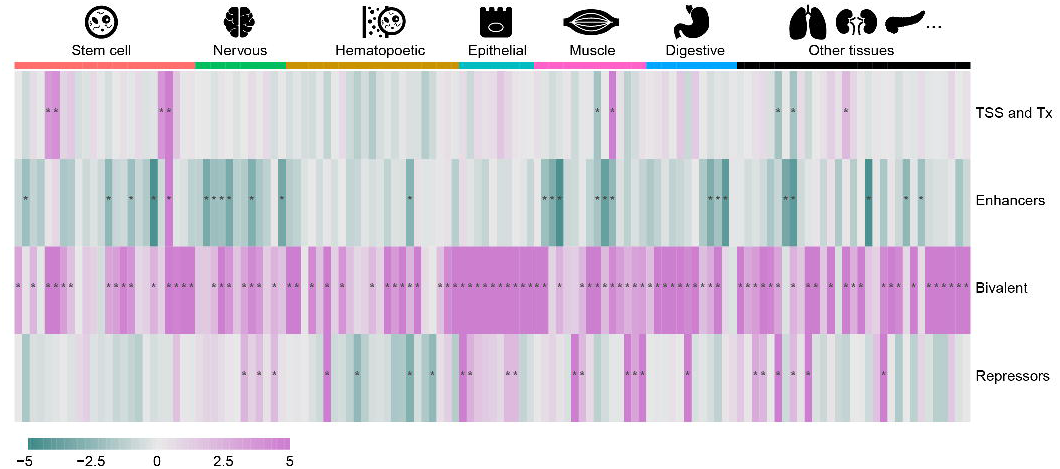
Enrichment and depletion of cxSVs in 127 Roadmap Reference Epigenomes across different chromatin states. Purple and green rectangles represent enrichment and depletion respectively, and deeper color represents smaller p-value. Significant enrichment or depletion (FDR <0.05) are marked by asterisks.

The function of bivalent chromatin states, which contain both the H3K27me3 repressive mark and the active H3K4me3 promoter mark and/or the active H3K4me1 enhancer mark, is proposed to keep environmentally and developmentally responsive genes in a poised state allowing for precise activation and silencing (Bernstein et al. 2006; Macrae et al. 2022). Key environmental factors that drive human adaptation to different geographical regions include infectious agents, diet, and climate (Hollox et al. 2022). Here, we find that the highest proportion of significant bivalent chromatin-state enrichments are for epithelial, digestive, and muscle reference epigenomes (Figure 5, Table S18). Since bivalent chromatin states associate with genomic regions that are active in a localized and temporal manner (Bernstein et al. 2006; Zhao et al. 2021), our results suggest that cxSVs are enriched in genomic regions that function in response to specific developmental or environmental factors.

### Rare cxSVs are enriched in neural genes

We next investigated the gene associations of common and rare cxSVs. We used FUMA (Watanabe et al. 2017) to perform gene ontology (GO) enrichment analysis for rare cxSVs (< 0.1% AF) in comparison against those that are common (≥ 0.1% AF). Of those 7,116 rare cxSVs, 3,304 overlap 2,897 unique protein-coding genes (Table S15) for which we find significant enrichment for those of neural functions (Figure 6c). After multiple test correction with FDR < 0.01, rare cxSVs are significantly enriched in 94 GO terms including synapse (q = 7.8e-9), neuron projection (q = 5.9e-7), synaptic membrane (q = 5.4e-9), and postsynapse (q = 5.2e-9) in the top 15 (Table S19). In contrast, common cxSVs are significantly enriched in only 9 GO terms (Table S20) with one for GO Cellular Component (Figure 6d) and with “nervous system process” being the least significantly enriched (q = 0.007) (Table S20). Enrichment fold change is significantly larger in rare cxSVs than common ones (binomial test, p = 0.0179, Figure S10). Our results suggest that cxSVs that are rare in the population or individually private disproportionately affect genes of neural function compared to common cxSVs. This suggests that there is a significantly high tendency for cxSVs to form in neural genes and a selective pressure against these cxSVs from rising in frequency in humans.

**Figure 6.**
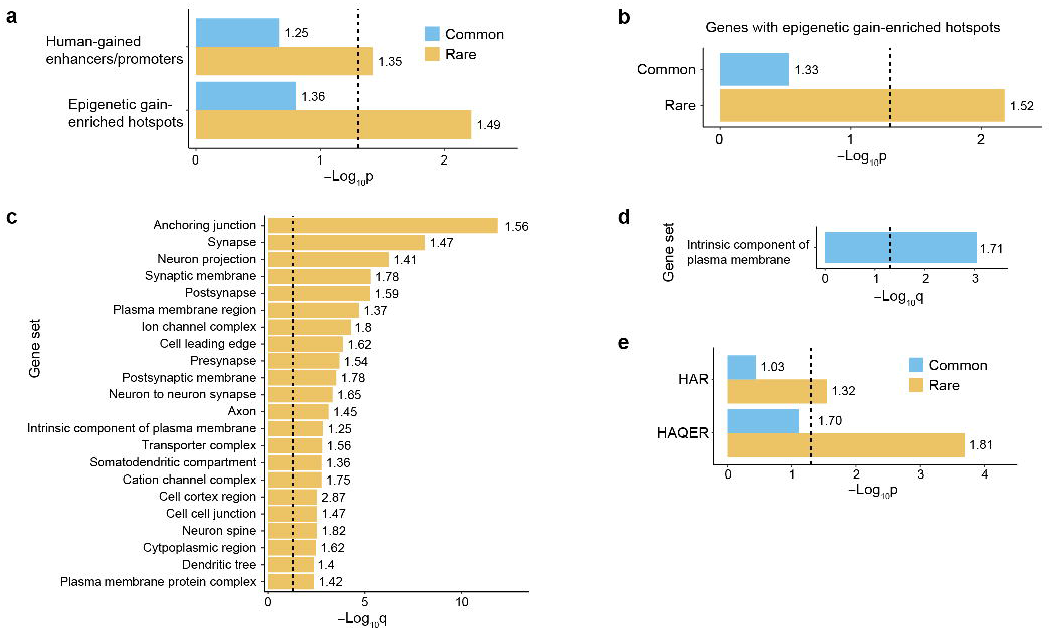
Enrichment of common and rare cxSVs in (a) “human-gained” enhancers/promoters, epigenetic gain-enriched hotspots, (b) protein-coding genes within epigenetic gain-enriched hotspots, (c-d) GO Cellular Component terms (with adjusted p-value), (e) HARs, and HAQERs.

### Rare cxSVs are enriched in genomic regions that recently diverged in humans

Many studies have identified rapidly evolving sequence changes in humans that are likely adaptive (Doan et al. 2016; Whalen and Pollard 2022; Mangan et al. 2022). Intriguingly, like cxSVs, many of these rapidly evolving regions are enriched at loci with high mutation rates (Mangan et al. 2022). We examined whether cxSVs might be occurring in genomic regions that are undergoing accelerated evolution in the human lineage and might contribute to ongoing adaptation in modern human populations. First, we investigated the co-localization between cxSVs and Human Accelerated Regions (HARs) (Doan et al. 2016), which are regions of the genome that are highly conserved across vertebrates but undergo accelerated sequence changes in the human lineage following human-chimpanzee divergence (Pollard et al. 2006). We find 58 HARs across 52 unique cxSV regions (Table S21) with significant overlap enrichment for rare cxSVs (permutation test, enrichment = 1.32, p = 0.028) but not for common cxSVs (Figure 6e), suggesting that in general, further changes to HARs may be selected against in modern humans. This is consistent with prior findings that sequence changes within HARs can be deleterious (Doan et al. 2016; Xu et al. 2015).

While HARs are regions that reflect human-specific adaptive changes of existing functional elements (Whalen and Pollard 2022), a recent study also identified functional genomic elements where the ancestral sequences were previously neutral, called “human ancestor quickly evolved regions” (HAQERs). HAQERs are the fastest-evolving regions in the human genome that rapidly diverged in the hominin lineage due to positive selection (Mangan et al. 2022). We find that 48/1581 unique HAQERs reside within 48 unique cxSV regions (Table S22). Like HARs, we find significant enrichment for HAQERs within rare cxSVs (permutation test, enrichment = 1.81, p = 0.00020) but not within common cxSVs.

Given that rare cxSVs are enriched in neural genes (Figure 6c, Table S19), we also examined whether cxSVs may affect neural enhancers/promoters in the developing cerebral cortex that recently evolved to have higher activity in humans (called “human-gained”) (Reilly et al. 2015). We find that while common cxSVs are not enriched (permutation test, p > 0.05), rare (AF < 0.1%) cxSVs are significantly enriched for human-gained enhancer/promoter regions (Figure 6a). Rare cxSVs are especially enriched at loci with a large number of human-gained enhancer/promoter regions, called “epigenetic gain-enriched hotspots” (permutation test, enrichment = 1.49, p = 1.3e-2) (Figure 6a). Furthermore, rare cxSVs (p = 6.7e-3) but not common cxSVs (AF ≥ 0.1%) (p = 0.33) are significantly enriched within the 152 protein-coding genes within these “epigenetic gain-enriched hotspots” (Figure 6b). These genes perform functions related to regionalization, neuron differentiation/development, and A/P pattern formation (Reilly et al. 2015). The enrichment of rare cxSVs but not common cxSVs at HARs, HAQERs, and human-gained enhancers/promoters suggest that there is a significantly high tendency for cxSVs to arise in these human-evolved regions and selective pressure against them rising in allele frequency.

However, there are a number of cxSVs common to specific populations that affect human-evolved regions and may contribute to ongoing adaptation. For example, a cxSV (chr9:13734361-13755897, ABC’B’D) that is at 0.18% AF in EAS but rare elsewhere overlaps two HARs (HAR34 and 2xHAR.19) (Figure 7a, Table S21). HAR34 was experimentally validated to be an active enhancer in facial mesenchyme and forebrain using a transgenic mouse model (Visel et al. 2007; Whalen and Pollard 2022), and 2xHAR.19 was demonstrated experimentally to be an active enhancer in neural progenitor cells (Uebbing et al. 2021). The right breakpoint of the inversion part of this cxSV breaks 2xHAR.19 (chr9:13755747-13756163), and concomitantly, the inversion also places HAR34 ∼9.5 kb away from its original genomic location (Table S21). Another example is cxSV on chr3: 2263830-2271090 (ADCDE) which is polymorphic across all continental superpopulations except OCN (Table S15) and co-localizes with HAR HACNS_888. This cxSV also overlaps a CTCF binding site (ENCODE accession E2174027), and both HACNS_888 and this CTCF binding site are within *CNTN4*, is a risk gene for neuropsychiatric disorders that modulates synaptic plasticity in the hippocampus (Oguro-Ando et al. 2021).

**Figure 7.**
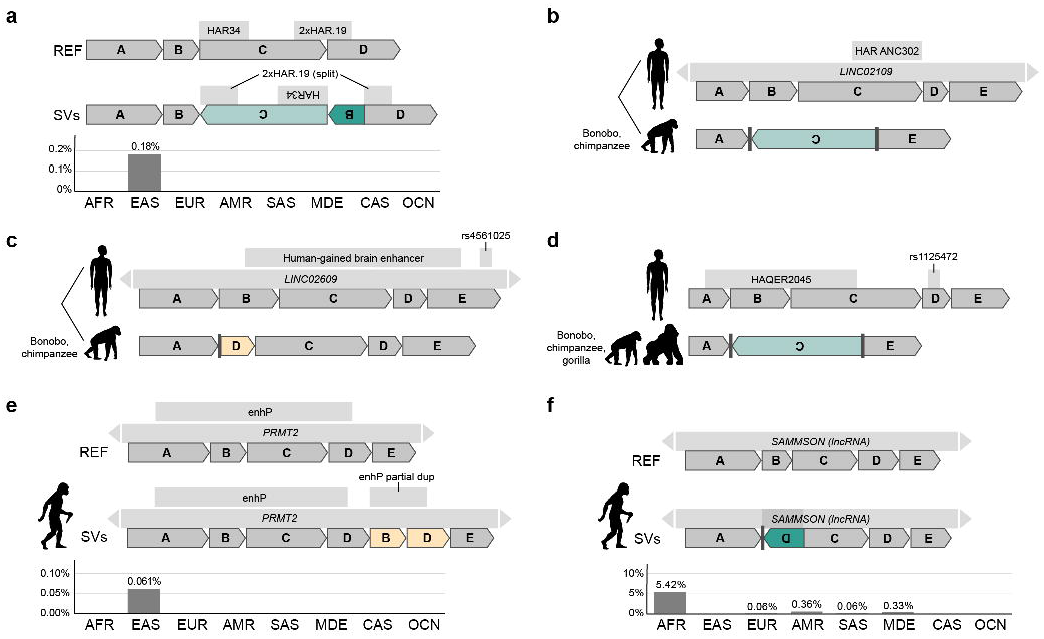
Examples of cxSVs at loci that recently diverged in humans. (a) cxSV identified in EAS only (0.18% AF) that affect HARs that are experimentally validated enhancers. Fixed loci in humans that diverged with bonobos and chimpanzees via a cxSV (with respect to hg38) (b) affecting HAR ANC302 within *LINC02109* and (c) within *LINC02609* that contain a human-gained brain enhancer and GWAS SNP rs4561025. (d) Fixed loci in humans that diverged with bonobos, chimpanzees, and gorillas affecting HAQER2045 and containing GWAS SNP rs1125472. (e,f) cxSVs identified in Neanderthal that are shared with humans within *PRMT2* and lincRNA *SAMMSON*. AF (%) indicated by bar-plot across superpopulations. ENCODE cCRE with proximal enhancer-like signature indicated by “enhP”.

We also identify a cxSV (ch9:110389915-110390324, ABBCDED’F) that disrupts HAQER0272 (chr9:110389548-110390242) (Table S22); both are located within a human-specific lincRNA *ENSG00000235281* with restricted testis expression (Baran et al. 2015). This cxSV has AF 0.1-0.3% in AFR, EUR, and SAS populations and is rare elsewhere (Table S15,S22). Another example is cxSV (chr5:26793060-26793640, ABCB’E), which disrupts HAQER0188 along with NANOG and SPI1 developmental transcriptional factor binding sites in the same locus (Hammal et al. 2022). This cxSV is common (AF 0.3%) only in the EUR superpopulation (Table S15,S22). Lastly, cxSV (chr1:237402804-237402905 AC’D), which is polymorphic (AF 0.3%) in the MDE population, disrupts HAQER0130 and the inversion of Block C breaks a binding site of PAX8 (Hammal et al. 2022) (Table S22), which is a thyroid developmental transcription factor (Fernández et al. 2015).

### Divergence between humans and non-human primates via cxSV

ARC-SV can also be used to identify cxSVs that differ between species. To investigate how cxSVs might be involved in the genome divergence of modern humans and our closest primate relative, we obtained paired-end whole-genome sequencing datasets for bonobos and performed cxSV detection with respect to GRCh38 and alignment-based SV validation against bonobo assembly (Mao et al. 2021). Overall, we find 377 loci that diverged between bonobos and modern humans via complex SVs that appear to be fixed within humans and derived within humans relative to other primates (Table S23, Table S15, Source Data). For example, we find a bonobo cxSV (chr5:29094936-29109104, AC’E) that encompasses HAR ANC302 (chr5:29105324-29105442) (Figure 7b). This 14 kb cxSV was orthogonally validated using PacBio long-reads of the same bonobo genome and against the bonobo genome assembly GCF_013052645.1 (Mao et al. 2021) (BLAT identity 98.6%) (Kuhn et al. 2013). It spans *LINC02109* which is a human-specific RNA expressed specifically in the testis (Baran et al. 2015). As this locus is fixed in the human population, we speculate that divergence from bonobos via this cxSV allowed for the rapid accelerated evolution of this locus in the human lineage, possibly through creating *LINC02109*. Another cxSV that diverged between bonobos and humans but is fixed within humans is within *LINC02609*, where the bonobo sequence structure is ADCDE at chr1:90794594-90794971. Intriguingly, this locus in the human genome contains a human-gained brain enhancer (chr1:90794843-90796293) (Reilly et al. 2015) and a GWAS-SNP rs4561025, which is associated with neuroticism (Cai et al. 2020) (Figure 7c). In addition, we also find a cxSV in bonobos relative to humans on chr11:92193737-92196296 (AC’E) where the cxSV overlaps HAQER0245 and is linked to GWAS-SNP rs1125472, which is associated with mathematical ability (Lee et al. 2018) (Figure 7d). This cxSV is also conserved in chimpanzees and gorillas i.e. only the human version of this locus is non-ancestral suggesting that this is a human-derived sequence that went to fixation after divergence from the most recent common ancestor (MRCA) with chimpanzees and bonobos. Overall, we find 41 of such fixed human-derived sequences where divergence with our MRCA is via a cxSV (Table S24). These findings suggest that cxSVs may play a role in the divergence of humans from other primates.

### cxSVs shared with Neanderthals

We also used ARC-SV to examine whether cxSVs that are polymorphic within humans are also identified in Neanderthals. In the Neanderthal genome (Prüfer et al. 2014), we discovered 5 cxSVs that are shared with modern humans (Table S25); 4 of which are common in the human population and validated in individuals with HPRC assemblies (Liao et al. 2022) (Table S3), and 1 of which is rare. One of these validated cxSVs in the Neanderthal genome is a dispersed duplication deletion (chr3:70399918-70401539, ADCDE, Figure 7f) that overlaps the oncogene *SAMMSON*, a lincRNA that is tissue-specifically expressed in more than 90% of human melanomas (Verfaillie et al. 2015). *SAMMSON* knockdown drastically decreases melanoma cell viability irrespective of *BRAF, NRAS*, or *TP53* mutations, whereas exogenous *SAMMSON* increases clonogenic potential (Leucci et al. 2016). This cxSV is highly prevalent in AFR populations (5.4% AF) and common in AMR (AF 0.36%) and MDE populations (AF 0.33%) populations but rare elsewhere (Table S15,S25). Because of its high prevalence in AFR populations, this cxSV allele may have been introduced into the Neanderthal genome by modern humans, possibly when making a failed migration from Africa to the Middle East more than 100,000 years ago (Chen et al. 2020). Another example is a cxSV allele shared only between Neanderthal and EAS populations (AF = 0.061%), suggesting the possibility of Neanderthal introgression. This shared cxSV disrupts a cCRE with proximal enhancer-like signature (E2148159) (The ENCODE Project Consortium 2012) and resides within *PRMT2*, which regulates glioblastoma (Dong et al. 2018; Liu et al. 2022) (Figure 7e, Table S25). Introgressed cxSVs from archaic humans may contribute to the adaptation and phenotypic variability seen in specific modern human populations (Huerta-Sánchez et al. 2014; Simonti et al. 2016; Vernot et al. 2016; Hollox et al. 2022).

## DISCUSSION

Here, we developed an innovative probabilistic model and machine learning framework ARC-SV to detect complex structural variation accurately, comprehensively, and automatically from standard WGS data that can also be implemented efficiently at population-scale. Importantly, ARC-SV operates using a “bottoms-up” approach where it does not look for data signatures to match pre-conceived structures (Collins et al. 2020; Byrska-Bishop et al. 2022) but rather the structures detected are purely data-driven (Supplementary Methods). Typical SV detection methods rely solely on the analysis of aberrant read signatures either of a single or multiple types (Escaramís et al. 2015; Lin et al. 2015; Guan and Sung 2016; Kosugi et al. 2019). In ARC-SV, we use such techniques to propose SVs, but ultimately combines the evidence both for and against each candidate in a holistic and more statistically rigorous fashion than previous methods.

By implementing ARC-SV to analyze 4,262 human genomes across the world, we show that complex genome structural variation is a major component of natural human genetic variation and pervasive in all human populations where individual superpopulations carry their own distinct cxSV signatures (Figure 3f-g, Table S15). By performing cxSV analysis between bonobos, the Altai Neanderthal (Prufer et al. 2014), and humans, we show that cxSVs may have contributed to divergence in the human lineage and identify cxSVs that may have entered the human population via Neanderthal introgression. We believe that future applications of ARC-SV to examine disease cohorts or differences between additional species will uncover novel aspects of disease and evolutionary biology.

cxSVs have a significantly high tendency to be found in regions with elevated *de novo* mutation rates and recombination initiation hotspots (Pratto et al. 2014; Zhao et al. 2020) (Figure 4a). Loci with elevated mutation rates likely contribute an excess of alleles for evolution (Xie et al., 2019; Mangan et al. 2022). Considering this, it is perhaps not surprising that rare cxSVs are significantly enriched in genomic regions undergoing rapid or accelerated positive selection in humans, i.e. HARs (Whalen and Pollard 2022; Doan et al. 2016) and HAQERs (Mangan et al. 2022). Although common cxSVs are not enriched for HARs and HAQERs, suggesting that most changes in these regions are likely to be deleterious, the over-representation of rare cxSVs in HARs and HAQERs suggests that these regions continue to be highly mutable and may serve as potential substrates for ongoing adaptation within humans. We have identified a number of common cxSVs overlapping HARs and HAQERs that have risen in frequency in modern humans and may underlie variation in the facial mesenchyme, neural tissues, testis, and thyroid. Intriguingly, the tissues with bivalent chromatin regions most significantly enriched for both common and rare cxSVs include epithelial, digestive, and muscle tissue (Figure 5), which parallel known adaptive changes to different environments, such as changes to climate, infectious agents, diet, and terrain (Hollox et al. 2022).

The other side of the coin for hypermutable genomic regions is of course their tendency to generate disease predisposing alleles. Although cxSVs are often omitted in current GWAS due to the limited ability to analyze SVs (Winkler et al. 2014; Torkamaneh and Belzile 2022), we find that the GWAS SNPs that most closely associate with cxSVs also associate with disease phenotypes in tissues that mark human-specific adaptations such as spine, gastrointestinal, and neural (Figure 4d, Table S16,S17). Interestingly, we find that cxSVs that are rare in the population or are private to individuals are enriched in genes with neurodevelopmental functions (Table S19). It is very plausible that leveraging ARC-SV to identify cxSVs in disease populations will reveal new disease-associated alleles and help uncover the the molecular pathophysiology of neurodevelopmental and neuropsychiatric disorders.

However, like any reference-based method using short-read, short-insert, WGS data, ARC-SV is also subject to uncertainty and lack of long of long-range information in short-read alignments. Currently, we show that ARC-SV outperforms long read-based cxSV detection methods such as SVision (Lin et al. 2022) (Table S13). With at least three new ultra-high-throughput short-read sequencers from three different companies just released this year to compete with the Illumina short-read platform (Pennisi 2022) and to meet the demand of new initiatives to sequence millions of individuals for national genome projects and population-scale studies (Wonkam 2021; Gaziano et al. 2016; Saunders et al. 2019), there remains a vital need to develop effective analysis methods such as ARC-SV to extract new dimensions of genetic information from short-read sequencing data. In anticipation of this new era of an unprecedented scale in the generation of population-level WGS data, we also show here that ARC-SV can be readily implemented for such competing short-read platforms Element Bioscience (Arslan et al. 2022) and MGI (Shen et al. 2020) where concordance for cxSV calls with Illumina data are 92% and 80% respectively (Table S26,S27).

A key component to the development of our statistical framework is leveraging the high-quality state-of-the-art diploid genome assemblies of 42 diverse individuals by the HPRC (Liao et al. 2022; Miga and Wang 2021) for building our machine learning models to predict cxSV events at a much greater population-scale. We anticipate that the performance of these models will substantially increase when we can re-train them with the upcoming release of the full assemblies of all 350 individuals (Liao et al. 2022; Miga and Wang 2021). Our method to automatically detect and accurately resolve cxSVs from standard paired-end WGS opens new doorways in future research, particularly for large cohorts, to address how such aspects of genome diversity can contribute to various biological traits, evolution, genome biology, and diseases such as cancer and neurodevelopmental disorders.

## Methods

### Datasets for population SV calling and validation

Data for population SV calling and analysis were obtained directly from https://www.internationalgenome.org. HG01783 was removed from analysis due to ambiguous ancestry curation (indicated as both AFR and EUR). HuRef and HepG2 WGS data were obtained from Zhou et al (Zhou et al. 2018a) and Zhou et al (Zhou et al. 2019) respectively. Alignment data for calling SVs to benchmarking against those that are manually curated from NA12878 were obtained from Zhou et al (Zhou et al. 2018b).

### SV detection

Complex and simple (deletions, tandem duplications, and inversions) SVs were called using both probabilistic and machine learning algorithm components (see Supplementary Methods for full description of probabilistic and machine learning models). Computation was performed using the Avitohol supercomputer (Atanassov et al. 2017). SVs with (1) greater than 85% overlap to simple repeat, low complexity, satellite repeat, and simple tandem repeat annotations (Kuhn et al. 2013), (2) greater than 50% overlap with segmental duplications, (3) affecting fewer than 50 bp (see Affected Sequence Calculation in Supplementary Methods), (4) overlap with SV blacklist regions (https://cf.10xgenomics.com/supp/genome/GRCh38/sv_blacklist.bed), (5) and overlap reference gaps were filtered out. Such SV filtering criteria was also used for paired-end WGS-based SV call comparisons. Filtering was implemented using the “arcsv merge-filter” function. Additionally, only SVs with split-read or paired-end read support ≥ 1 were retained.

### SV merging

For population SV calls, only high-confidence SV calls identified from machine learning prediction (Supplementary Methods) were retained. SVs at the same locus and of the same type were merged into a master list. cxSVs at the same locus (50% reciprocal base pair overlap) and of the same type were merged.

### Genomic DNA

Genomic DNA sample of Craig Venter (NS12911) was purchased from the Coriell Institute. HepG2 DNA was extracted from cells using Qiagen DNeasy Blood & Tissue Kit (Cat No. 69504), and concentration was quantified using the Qubit dsDNA BR Assay Kit (Invitrogen, Waltham, MA, USA). Purity of gDNA (OD260/280 > 1.8; OD260/230 > 1.5) was verified using NanoDrop (Thermo Scientific, Waltham, MA, USA).

### Experimental validation of complex SVs

PCR primers were using Primer 3 (http://bioinfo.ut.ee/primer3-0.4.0/primer3/) and verified using UCSC In-Silico PCR tool (http://genome.cse.ucsc.edu/cgi-bin/hgPcr). Each complex SV was sequence assembled with 150 bp of flanking genomic sequence on each side (Supplementary Table S7), and primers were designed to bind the flanking sequences and to amplify the assembled cxSV in its entirety including all predicted breakpoints. Each target was amplified using 50 ng of genomic DNA. The amplification was performed using Phusion Green high-fidelity DNA polymerase (Thermo Fisher Scientific Inc, Waltham, MA) under the following conditions: initial denature at 98°C for 30 sec, total 30 cycles of: 98°C for 10 sec, 60°C for 30 sec, and 72°C for 1 min (or 2 min for longer PCR product), and final extension at 72°C for 10 min. The PCR products were then analyzed by agarose gel electrophoresis. PCR products were then purified using Zymoclean Gel DNA Recovery Kit (Zymo Research, Irvine, CA) and submitted for Sanger sequencing at Sequetech Corporation (Mountain View, CA).

### AF calculation

Allele frequency (AF) is calculated as the number of cxSV events divided by the number of haplotypes (i.e. 2 x number of samples). AF is calculated within all samples across different populations. Common and rare cxSVs are defined as those whose AF is larger and smaller than 0.001, respectively.

### Principal component analysis (PCA)

We performed PCA for common cxSVs using R function “prcomp”. For the top two PCs, we identified the top ten cxSVs with largest positive loadings and the top ten cxSVs with largest negative loadings as the population-specific cxSVs. We extracted 63 protein coding genes whose distance to the nearest population-specific cxSV is smaller than 100KB to perform enrichment analysis (see below).

### gnomAD-SV LD maps

The Supplementary Table 7 of Collins et al (Collins et al. 2020) catalogued 15,634 SVs (AF ≥ 1%) in strong LD (R2 ≥ 0.8) with at least one SNV within +/- 1Mb on the basis of 3,470 AFR and 1,883 EUR samples from the gnomAD datasets. We used these publicly available LD maps to examine LD-distance relationship between gnomAD-SVs and SNVs, which further guided our choice of window size (125 kb) for mapping common SNPs to cxSVs in the present study.

### GWAS Catalog lookup

For each cxSV, we looked up GWAS SNPs within its +/- 125 kb interval in the GWAS Catalog (Sollis et al. 2023) version 1.0.2 mapped to genome assembly GRCh38.p13 and dbSNP Build 154 (https://www.ebi.ac.uk/gwas/docs/file-downloads). We only used GWAS SNPs reaching the genome-wide significant level (p ≤ 5e-8) in this analysis.

### Heritability enrichment

We quantified the heritability enrichment of common SNPs (MAF ≥ 0.05) within +/- 125 kb of cxSVs across 1087 GWAS datasets (with total h2 Z-score ≥ 2) as previously described (Zhu et al. 2022). In brief, we first created the binary annotation indicating if a common SNP falls inside +/- 125 kb of any cxSV, and then used S-LDSC (Finucane et al. 2015) to estimate the standardized effect size of the binary annotation (defined as the 1-standard deviation increase of the binary annotation) conditioning on 96 baseline genomic annotations (Hujoel et al. 2019) in each GWAS, and lastly computed a one-sided p-value testing if the standardized effect size > 0. Given a GWAS dataset, a significantly positive standardized effect size indicates a significant heritability enrichment. As the 1087 GWAS datasets were mapped to genome assembly GRCh37, we lifted over cxSVs from GRCh38 to GRCh37 using UCSC liftOver (Kuhn et al. 2013) in this analysis.

### SV enrichment analysis

We merged all unique cxSVs detected into non-overlapping genomic segments (n=7194) and examined if these genomic segments are associated with elevated recombination rates. We performed a permutation test to assess the significance of enrichment. For each permutation, we randomly selected 7194 non-overlapping regions from the genome with their sizes matching the cxSV regions, and calculated permuted overlaps defined as the number of regions that overlapped recombination hotspots (Pratto et al. 2014). The permutation procedure was repeated 100000 times. The enrichment was defined as the ratio between the data overlap (number of non-overlapping cxSV regions that overlap recombination hotspots) and the mean of 100000 permuted overlaps, and the significance p-value was calculated by comparing the data overlap with 100000 permuted overlaps. We also used the permutation test to investigate the enrichment of cxSVs in HARs, HAQERs, chromatin regions from four categories and 127 cell types, as well as human-gained enhancers. To investigate whether the cxSVs are prone to elevated mutation rate, we performed a permutation test to examine the enrichment of germline DNMs in the 7194 cxSV regions. In this specific analysis, data overlap and permuted overlap were defined as the number of DNM events falling into the 7194 cxSV regions and 7194 permuted regions, respectively. The enrichment and significance p-value were calculated similarly as in the enrichment analysis for recombination hotspots. Using GENCODE V42 (Frankish et al. 2021), we extracted 3130, 2897, and 566 genes that overlap all cxSVs, rare cxSVs, and common cxSVs, respectively from 19086 autosomal protein coding genes. Here the overlapped genes are defined as those whose distance to the nearest cxSV is less than 5 KB. Then we used hypergeometric test to examine the enrichment of these cxSV genes in the neurodevelopmental enhancer gene set as well as the gene ontology terms (Watanabe et al. 2017).

### SV detection in Neanderthal and bonobo genomes

Neanderthal WGS reads were obtained from https://www.eva.mpg.de/genetics/genome-projects/neandertal (Prüfer et al. 2014). Bonobo WGS reads were obtained from SRA (accession SRR13998205). WGS reads were aligned to hg38 (GCA_000001405.15) using BWA-MEM (Li and Durbin 2009) version 0.7.15-r1140 (parameters -Y -K 100000000). PCR duplicates were marked using Picard tools version 2.4.1 (http://broadinstitute.github.io/picard/). SV calls were called by ARC-SV (see SV detection section). For Neanderthal SV calls, only those with corresponding SV calls (95% reciprocal SV coordinate overlap and matching SV type) in modern human populations were retained. All bonobo SVs were sequence assembled with 150 bp of flanking sequence tagged on both ends and validated against bonobo genome assembly (GCA_013052645.3) using BLAT version 3.5 (Kuhn et al. 2013) requiring a minimum of 97.5% sequence identity, that query and target alignment range are within 2.5%, that total target and query gap bases are no more than 2.5% of total size, and that query alignment must initiate within the first 50 bp of 5’ tagged flanking sequence and continuously align/match target until at least the last 50 bp of the 3’ tagged flanking sequence. Same workflow was used for comparison against chimpanzee (GCF_002880755.1) and gorilla (GCF_008122165.1) genomes.

## Author contributions

A.E.U. and W.H.W. supervised the study. B.Z., J.G.A., H.G., and X.Z. wrote the manuscript. A.E.U., W.H.W., B.Z., D.P., X.Z., G.S., and H.P.J. contributed resources. J.G.A. and W.H.W. conceived and designed the statistical framework for SV detection. B.Z., T.K., and G.S. designed and implemented machine learning framework for SV detection. J.G.A. wrote the software package. B.Z. conceived and designed method validation and implementation across datasets. B.Z., Y.H., and R.P. performed experiments. B.Z., C.R.H., Y.H., D.P., and H.L. performed SV calling, validation, and comparative analysis. H.G. performed population SV analysis. X.Z. designed and performed GWAS catalogue and heritability enrichment analysis. H.G. designed and performed all enrichment analysis.

## Acknowledgments

We thank Dr. Douglas F. Levinson, Dr. Bernardo Rodriguez Martin, Dr. Heinrich Dohna, and Dr. Janet Song for manuscript proofreading and providing feedback.

B.Z. is funded by NIH grant K01MH129758 and the Stanford Maternal and Child Health Research Institute Instructor K Support Award. J.G.A. received funding from NIH training grant T32-GM096982 and NSF Graduate Fellowship DGE-114747. H.L. and H.P.J. were funded by NIH grant U01HG01096 and R01HG006137. G.S. is funded by the National Research Foundation of Korea (NRF) grant funded by the Korea government (MSIT) (No. 2021R1A2C2010775). D.P. acknowledges the provided access to the e-infrastructure of the NCHDC – part of the Bulgarian National Roadmap on RIs, with the financial support by Grant No D01-168/28.07.2022. X.Z. is supported by Stein Fellowship from Stanford University, Institute for Computational and Data Sciences Seed Grant and Consortium on Substance Use and Addiction Seed Grant from Pennsylvania State University. A.E.U. and W.H.W. are funded by NIH grant P50HG00773506 (renewal of P50HG007735). W.H.W is also funded by NIH grant R01HG010359 and NSF grant DMS1952386.

