## Supplementary Figures for "Automatic detection of complex structural genome variation across world populations"

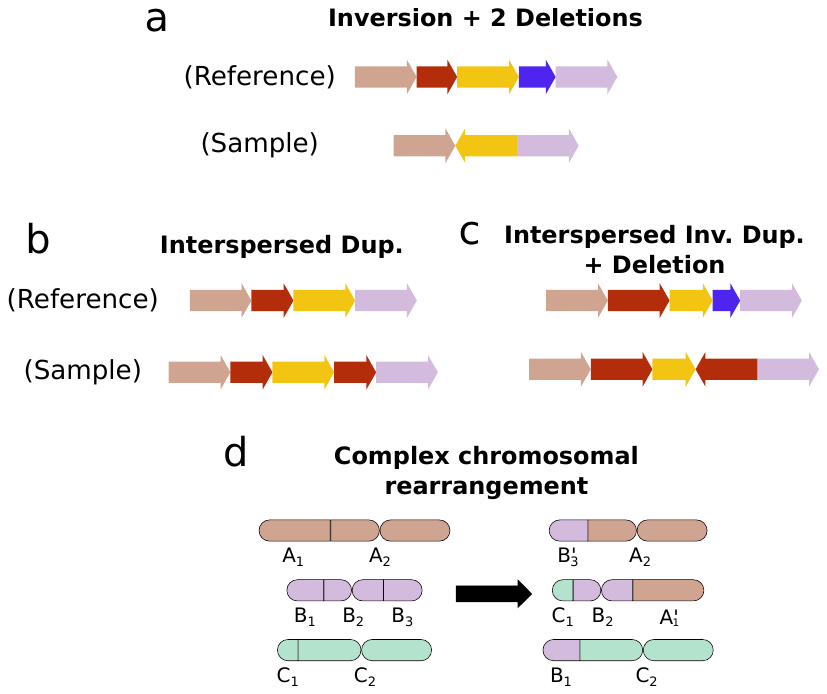

Supplementary Figure S1. Examples of complex SV structures. Examples (a), (b), and (c) fall under our category of localized cxSVs, but (d) does not.

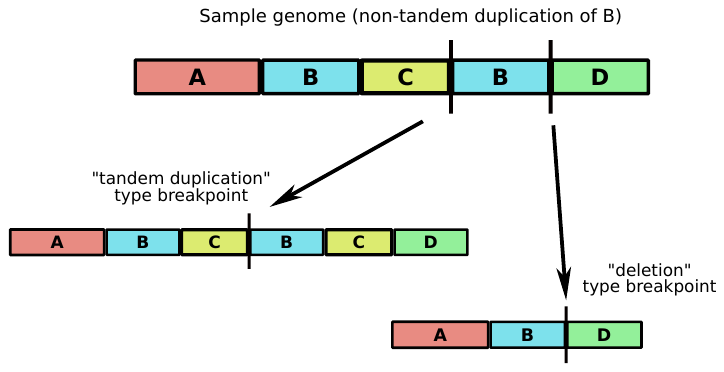

Supplementary Figure S2. Complex SVs generally introduce breakpoints that are consistent with multiple overlapping simple SVs. In this case, a non-tandem duplication of B (top) produces breakpoints consistent with a tandem duplication of BC (bottom left) and a deletion of C (bottom right). A naive SV detection method that equates breakpoints with SVs will thus produce two overlapping predictions, neither of which is correct.

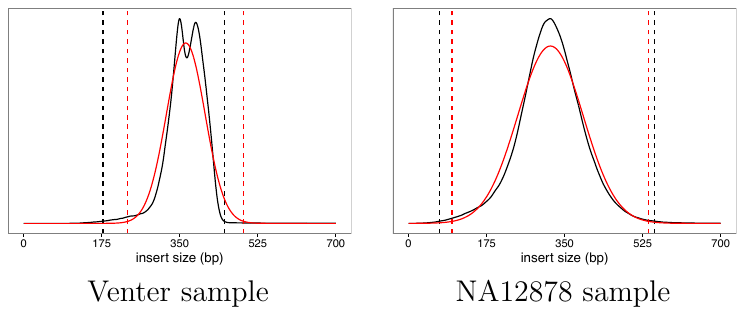

Supplementary Figure S3. Smoothed insert size distributions (solid curves) and insert size cutoffs used to define discordant read pairs. Our results based on a kernel density smooth and likelihood ratio statistic are in black, while the more typical normal approximates are in red.

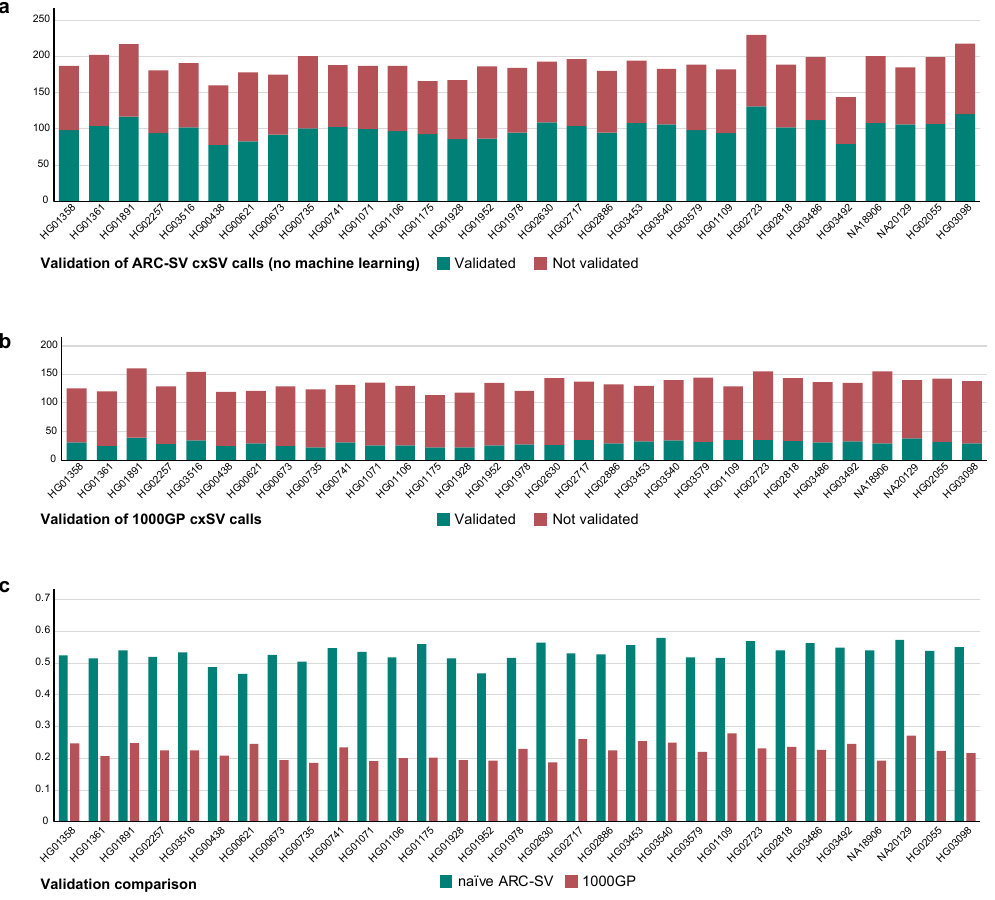

Supplementary Figure S4. Validation of cxSV calls in training genomes. (a) Alignment-based validation of count of (a) naïve ARC-SV cxSV calls and (b) 1000GP cxSV calls (Byrska-Bishop et al. 2022) in 31 training genomes for training machine learning models to predict SV validation. (c) Comparison of validation rate between naïve ARC-SV cxSV calls 1000GP cxSV calls.

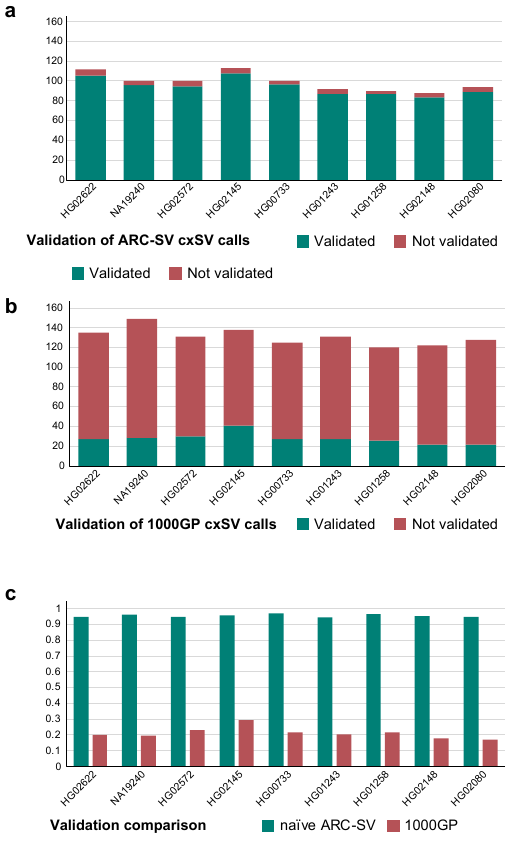

Supplementary Figure S5. Validation of cxSV calls in test genomes. (a) Alignment-based validation of count of (a) ARC-SV cxSV calls with implementation of machine learning model and (b) 1000GP cxSV calls (Byrska-Bishop et al. 2022) in 9 test genomes for training machine learning models to predict SV validation. (c) Comparison of validation rate between ARC-SV cxSV calls 1000GP cxSV calls.

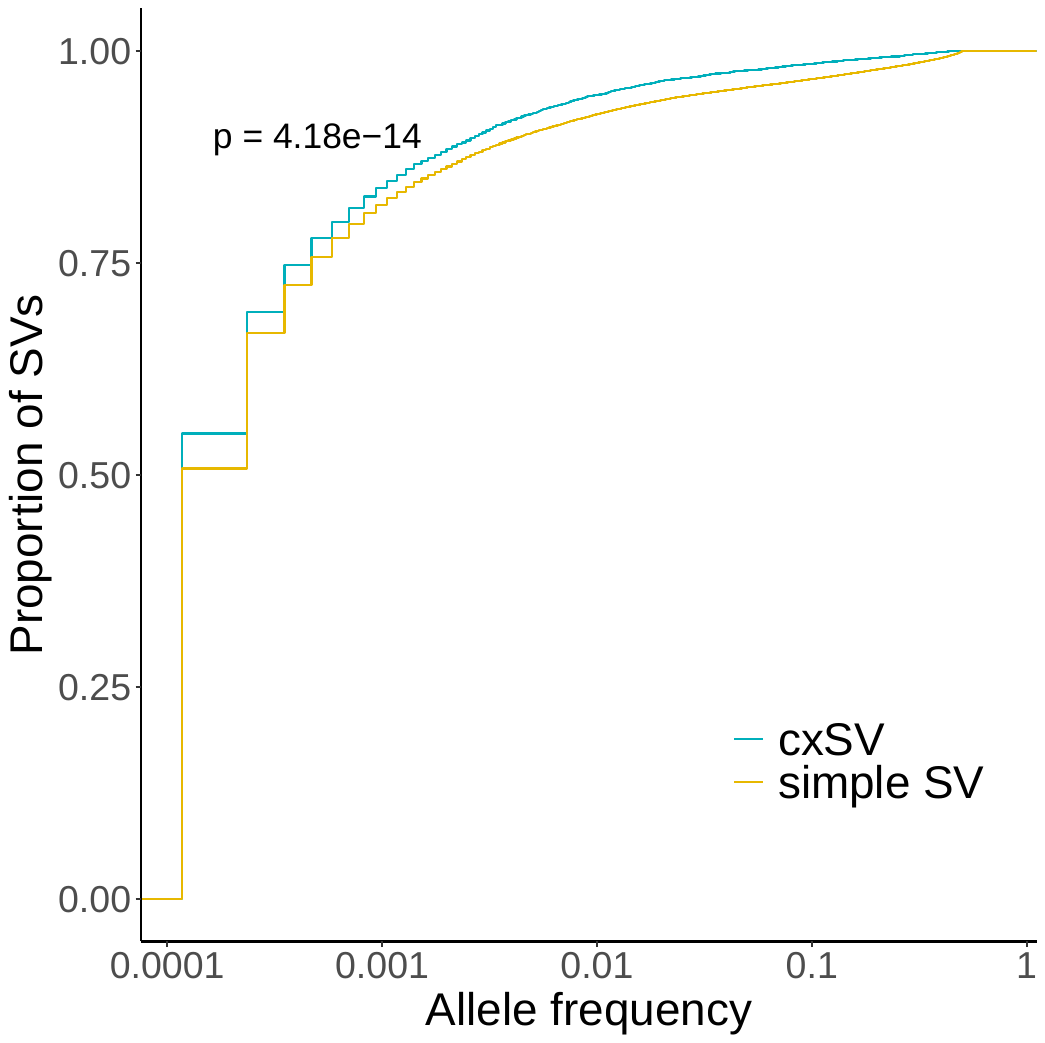

Supplementary Figure S6. Comparison of allele frequencies between simple and cxSVs. P-value for one-sided Mann-Whitney test.

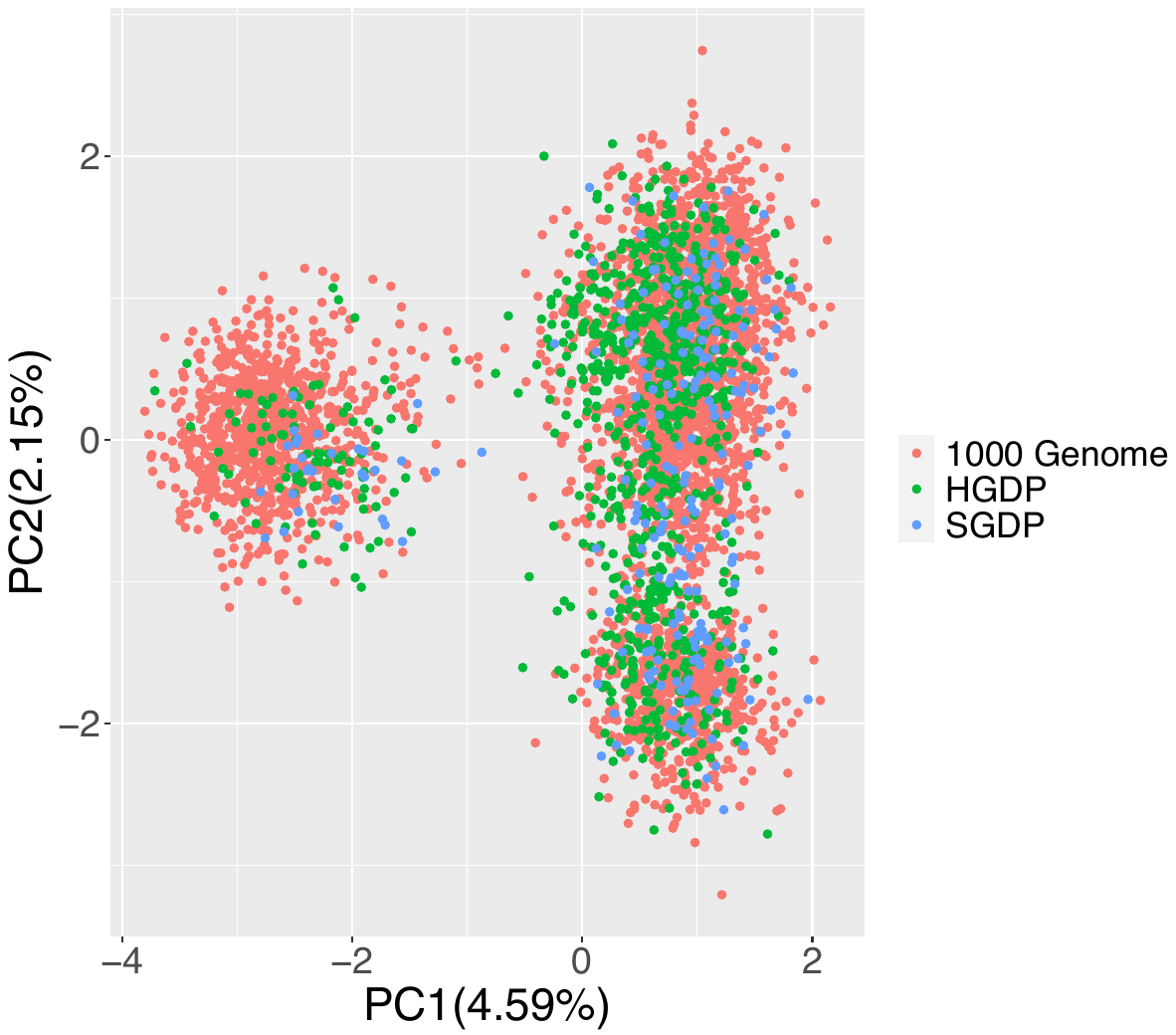

**Supplementary Figure S7**. PCA analysis of samples to ascertain batch effects of source datasets (1000 Genomes, Human Genome Diversity Project, Simons Genome Diversity Project).

Supplementary Figure S8. xSV regions that distinguish populations. Top two principal components (a) and (b) respectively that distinguish different populations. Top ten rows and bottom ten rows indicate different cxSV regions with top positive and top negative loadings respectively, and columns indicate different populations within superpopulations. Purple and gray colors indicate 1 (presence) and 0 (absence) respectively. (c) Enrichment in GWAS significant hits for genes located near the top cxSVs contributing to population-specific signatures.

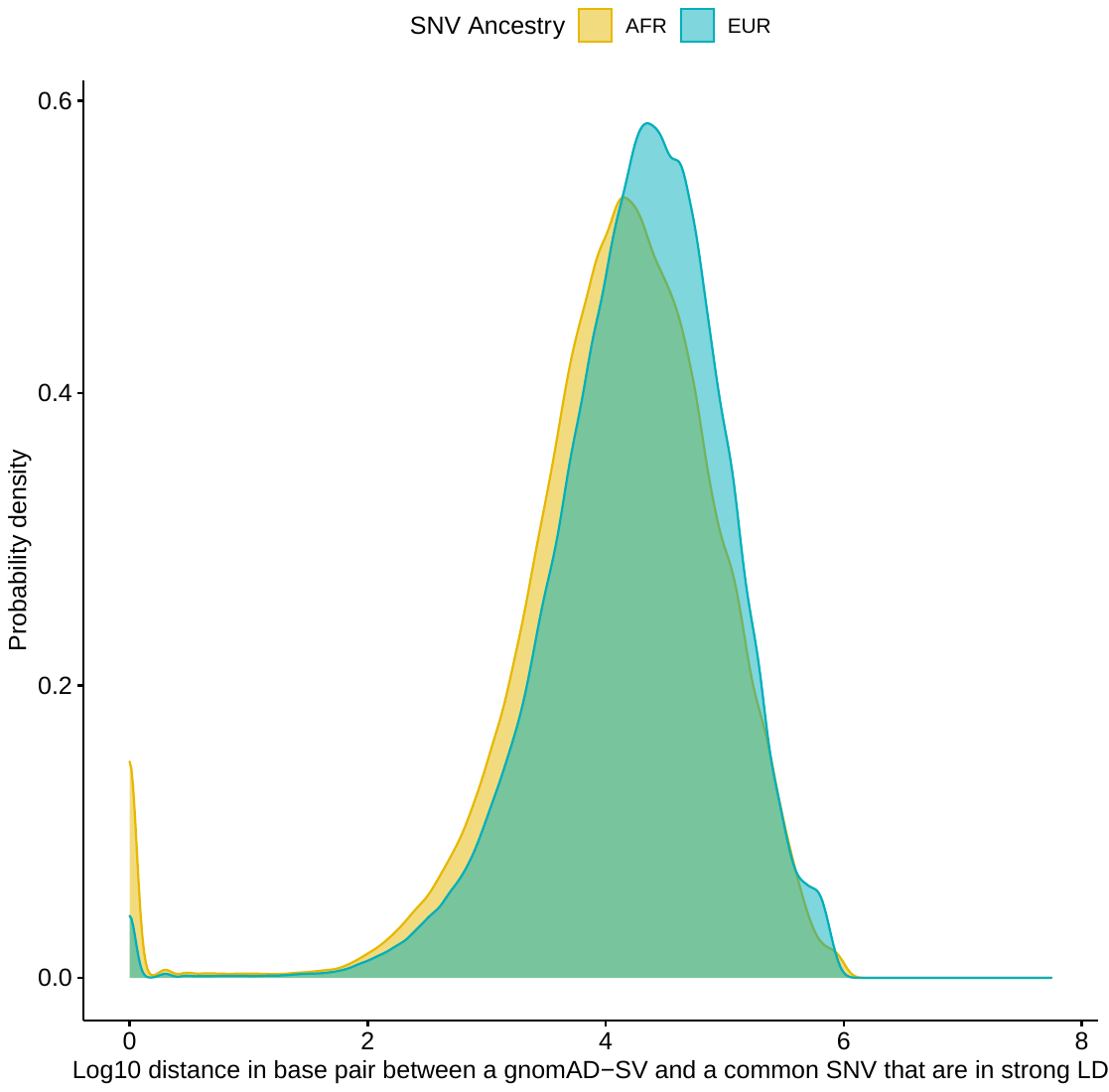

Supplementary Figure S9. Distance distribution of SNVs in strong LD with gnomAD-SVs (R2 ≥ 0.8), stratified by African and European ancestries.

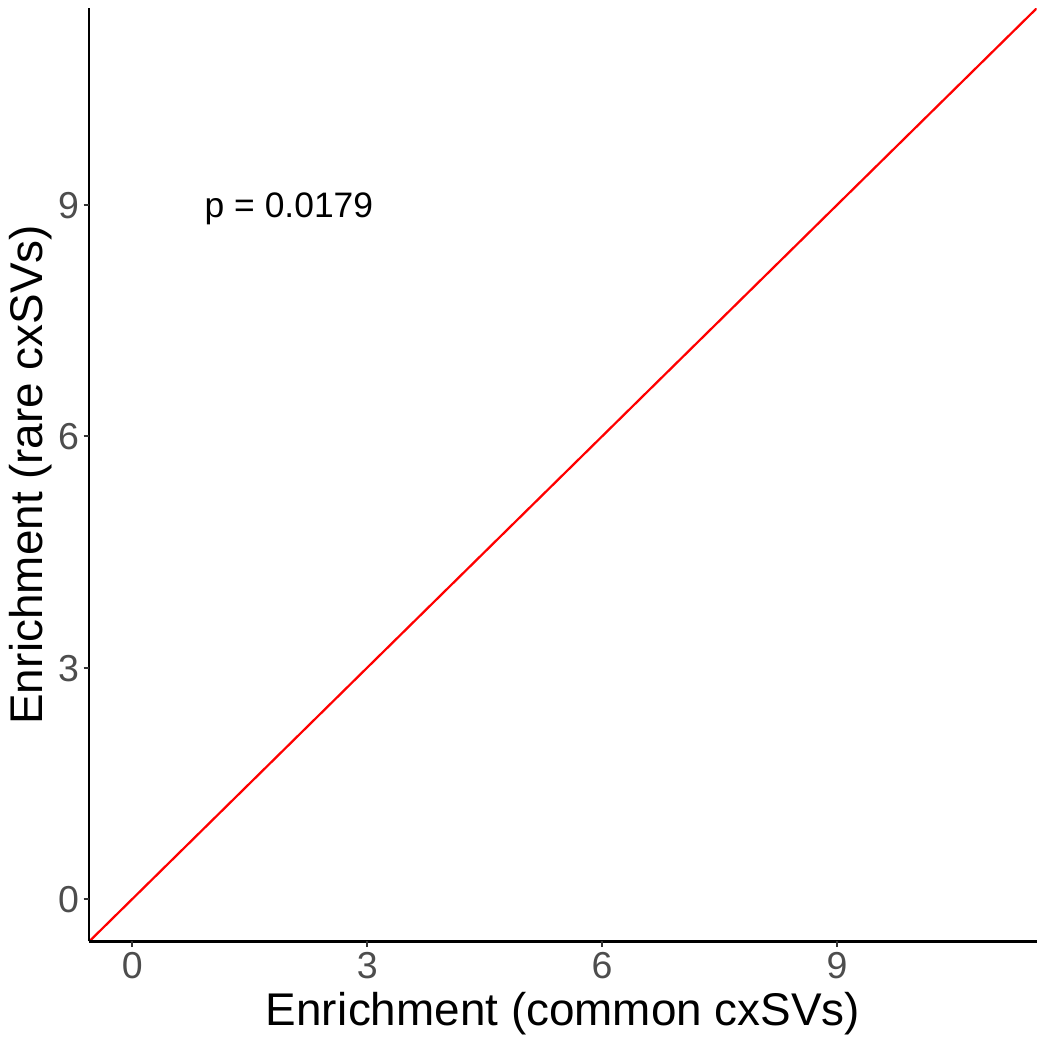

Supplementary Figure S10. Enrichment fold change is significantly larger for rare cxSVs than common cxSVs via binomial test.

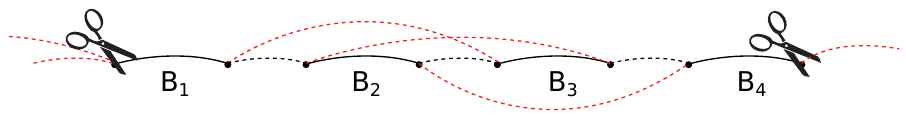

Supplementary Figure S11. Example of adjacency graph partitioning. The solid edges are block edges, and the dotted edges are adjacency edges. In this case, we are assuming there are no edges spanning the entire region shown (i.e., from before block B1 to after B4). The subgraph from B1 to B4 will be extracted, and alternating paths between B1 and B4 represent candidate SVs within the corresponding genomic region.

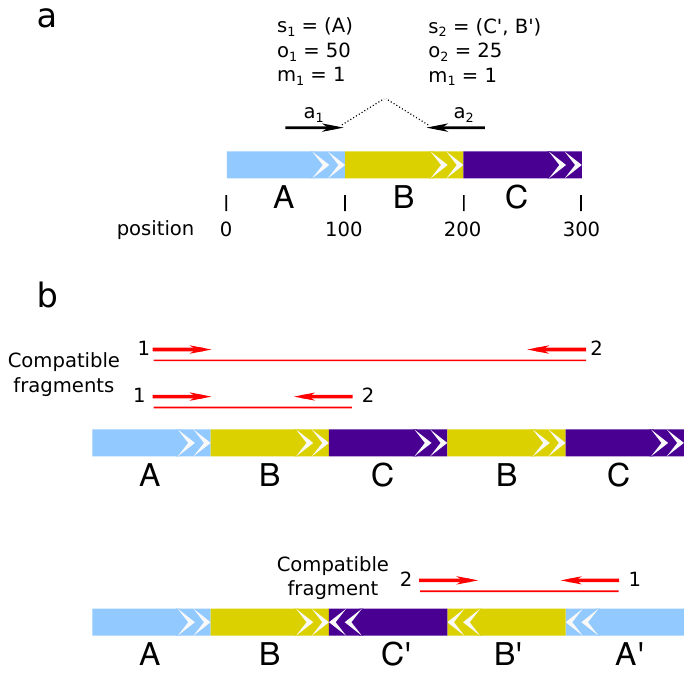

Supplementary Figure S12. Illustration of fragment model (a) and compatible fragments concept (b). The paired alignment $(a_{1},a_{2})$ has two compatible fragments under $\theta_{1}=ABCBC$ and one compatible fragment under $\theta_{2}=ABC^{'}B^{'}A^{'}$

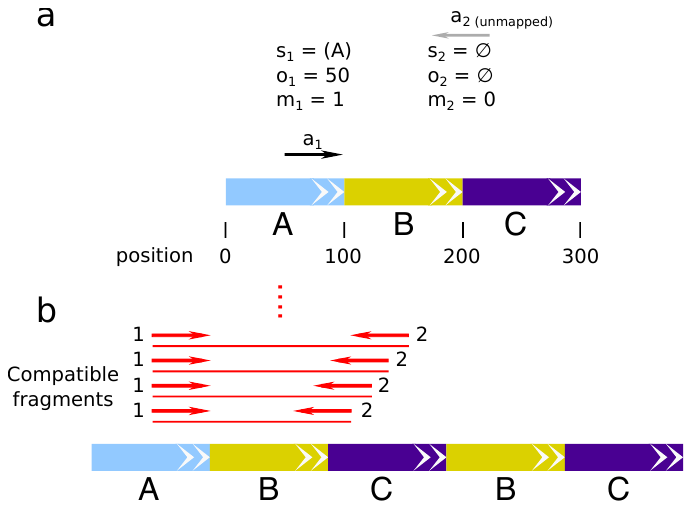

Supplementary Figure S13. Illustration of the alignment representation for hanging reads. (a) In the paired alignment $(a_{1},a_{2})$, read 2 is unmapped as indicated by $m_{2}=0$; (b) thus, any fragment matching $a_{1}$ is compatible.

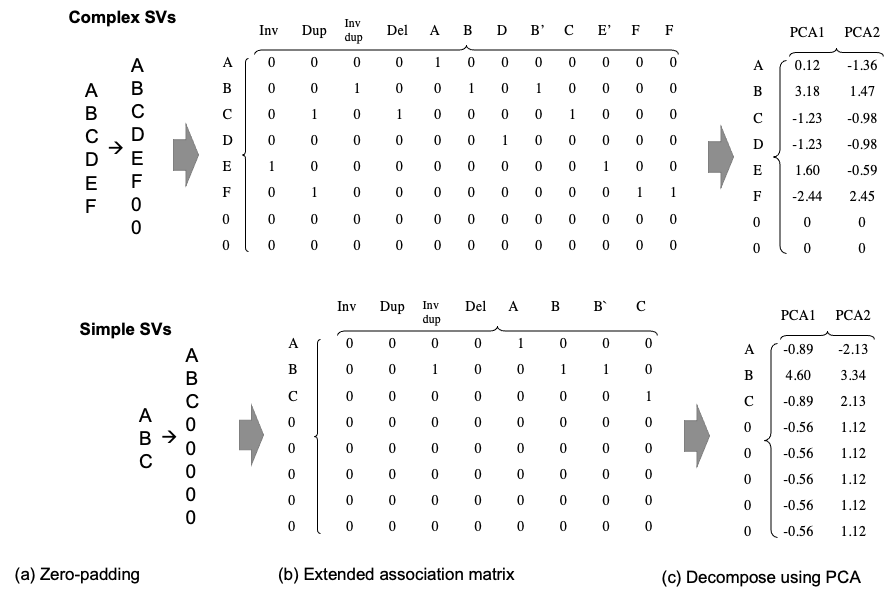

Supplementary Figure S14. Illustration of matrix transformation of reference and rearrangement block configurations of complex and simple SVs for machine learning. (a) Representation of reference blocks with zero padding as a one-hot vector. (b) Extended mapping matrix of reference blocks and its rearranged blocks where “1” indicates SV event (Inv, Dup, INV dup, Del) occurrence. (c) Decomposition of matrix using PCA.
