## Supplementary Methods for "Automatic detection of complex structural genome variation across world populations"

**Detection of complex structural variation using probabilistic modeling**

***Notation***

Intervals. Genomic positions are discrete thus any intervals should be understood as to contain only integers, i.e., $\left[ a,b \right)=\{z\mathbb{\in Z :}a\leq z<b\}$.

Genomic blocks. We refer to a sequence of genomic segments or “blocks” using notation such as ABCDE, where A denotes the first block, B the second, etc., and where AB’CDE denotes that block B is inverted (reverse complemented).

***Breakpoint discovery***

Given a coordinate-sorted data set of paired-end read alignments, the first step of our procedure is to determine candidate breakpoints in the reference genome, i.e., positions that are the sites of novel adjacencies in the sample genome.

1. **Breakpoint discovery by soft-clipped and split alignments**

Soft-clipped and split alignments from coordinate-sorted paired-end read alignments are used for breakpoint detection. Split alignments (MAPQ score ≥ 20) are assigned (1) a type (deletion, duplication, left/right side of inversion) and (2) a breakpoint interval based on the orientation and mapped positions of the primary and secondary alignment. For soft-clipped reads without split alignments, the soft-clipped portion is required to be ≥ 5 bp, to have a median base quality ≥ 15 and a MAPQ score of ≥ 10, for increased sensitivity.

Soft-clipped reads within 5 bp and with the same clip orientations are merged; the breakpoint position whose supporting reads have the highest total MAPQ is used as a candidate. As sequence microhomology can lead to two soft-clip clusters at a single insertion or inversion breakpoint, to avoid creating the artefact of two breakpoints, each pair of soft-clip clusters with overlapping aligned bases and opposite clip orientations are merged whenever the two breakpoints are within 25 bp; the region between the breakpoints is then used as a breakpoint interval.

2. **Breakpoint discovery by discordant read pairs**

Discordant read pairs (DRPs) are abnormal paired-end alignments that suggest the presence of structural variation (SV) where abnormal insert sizes suggest deletions and insertions and abnormal mapping orientations suggest tandem duplications and inversions. DRPs from the left and right sides of inversions are treated separately since they indicate separate breakpoint locations. Each of these 5 types of DRPs is handled separately for the purpose of breakpoint detection.

Deletion and insertion DRPs are identified according to insert size cutoffs, so first the density of the insert size distribution $f$ must be estimated. This is done by sampling properly oriented read pairs (1 million by default) from randomly selected genomic regions and applying kernel density estimation to the observed (mapped) insert sizes.

Conventionally, discordant insert sizes are those outside of $(\mu-n\sigma,\mu+n\sigma)$, where $\mu$ and $\sigma$ are the mean and standard deviation of the estimated insert size distribution $f$, and the multiplier $n$ is commonly set to 3 (Layer et al. 2014; Rausch et al. 2012; Yalcin et al. 2012). To allow for a variety of different insert-size distributions, cutoffs based on a likelihood ratio is used instead. We define deletion pairs as those with insert sizes exceeding $z^{*}$, where $z^{*}$ is the smallest value such that, for all $z>z^{*}$,

$$\frac{\max_{S\geq0}f(z-S)}{f(z)}>C.$$

The denominator above is just the likelihood of an insert size z if the sample genome matches the reference. The numerator is obtained by maximizing the likelihood over all deletion sizes, i.e., over all upward shifts in the insert size distribution. We set $C=\exp(3^{2}/2)$ so that, if $f$ is a normal distribution, the cutoff $z^{*}$ is the commonly-used value $\mu+3\sigma$ (Layer et al. 2014; Rausch et al. 2012; Yalcin et al. 2012). Insertion read pairs are defined by insert sizes smaller than $z_{*}$, where $z_{*}$ is defined analogously to $z^{*}$ but with maximization over $\{S\leq0\}$. Examples from our data are shown in Supplementary Figure S3.

3. **Discordant read pair clustering**

For discordant pair $i$, we denote the mapped insert size as $z_{i}$ and the $3^{'}$-most mapped read positions as $a_{i}$ (leftmost read with respect to the reference genome) and $b_{i}$ (rightmost). After identifying DRPs, we construct a graph with nodes corresponding to DRPs. Two DRPs $i$ and $j$ are connected if they are of the same type, if both $|a_{i}-a_{j}|$ and $|b_{i}-b_{j}|$ are less than $\mu+3\sigma$, and if $\max\{a_{i},a_{j}\}<\min\{b_{i},b_{j}\}$. The third condition requires that some region is spanned by both DRPs. (For insertions, we require $\max\{a_{i},a_{j}\}-h<\min\{b_{i},b_{j}\}$ to allow for up to $h=20$ base pairs of microhomology at the insertion site.) The parameters $\mu$ and $\sigma$ are estimated from the insert size distribution using the median and IQR/1.35, respectively, where IQR is the interquartile range. After constructing the graph of compatible read pairs, we cluster the read pairs by taking the largest clique(s) in each connected component (ignoring singleton components).

4. **Discordant read pair FDR**

We select “significant” clusters using a procedure that adapts to the coverage of the data and only requires a target false discovery rate (FDR). For a given DRP type with $M$ total clusters, we assign each cluster $j$ a heuristic score $s_{j}$ indicating the strength of evidence. For inversions and tandem duplications, $s_{j}$ is the cluster size. For insertions and deletions, $s_{j}$ is a log-likelihood ratio statistic. For deletion clusters, we first estimate the deletion size ${\overset{̂}{D}}_{j}$:

$$D_{j}=\min\left\{ \frac{1}{|\mathcal{C}_{j}|}\sum_{i\in\mathcal{C}_{j}} z_{i}-\mu,\min_{i\in\mathcal{C}_{j}}b_{i}-\max_{i\in\mathcal{C}_{j}}a_{i} \right\},$$

where $\mathcal{C}_{j}$ is the set of read pairs within the cluster. The cluster score is

$$s_{j}=\prod_{i\in\mathcal{C}_{j}} \frac{\phi(z_{i};\mu+D_{j},\sigma)}{\phi(z_{i};\mu,\sigma)},$$

where $\phi(\cdot;\mu,\sigma)$ is the density function for the normal distribution. The case for insertions is completely analogous.

Under a null hypothesis of no SV, we would still expect some false DRP clusters to form when independent spurious discordant alignments occur close together. To simulate the formation of false clusters, we move each DRP to a new reference position uniformly at random, without changing the insert size or mapping orientations. Repeating the clustering process on these DRPs yields $M_{0}$ clusters. The resulting distribution of cluster scores is used as an estimate ${\overset{̂}{F}}_{0}$ of the null score distribution. We also use ${\overset{̂}{\pi}}_{0}=\min\{1,M_{0}/M\}$ to estimate the proportion of false clusters in the original sample. If FDR control at level $\alpha$ is desired, we use a cluster score cutoff

$$s^{*}=\min_{j}\left\{ s_{j} : \frac{M{\overset{̂}{\pi}}_{0}(1-{\overset{̂}{F}}_{0}(s_{j}))}{\#\{k:s_{k}\geq s_{j}\}}\leq\alpha\right\}.$$

The procedure above is very similar to that of Benjamini and Hochberg for general FDR control of multiple hypothesis tests (Benjamini and Hochberg 1995). Suppose $F_{0}$ is the (continuous) null distribution of some test statistic $s_{j}$. If the distribution of this statistic were stochastically larger under the alternatives of interest, one would use $1-F_{0}(s_{j})$ as a *p*-value. If there are $M$ independent test statistics $\{s_{1},\ldots,s_{M}\}$, then the Benjamini-Hochberg procedure rejects hypothesis $j$ if $s_{j}$ exceeds

$$s^{*}=\min_{j}\left\{ s_{j} : \frac{M(1-F_{0}(s_{j}))}{\#\{k:s_{k}\geq s_{j}\}}\leq\alpha\right\}.$$

where $\alpha$ is the target FDR. The left-hand side of the inequality above is an estimate of the false discovery proportion (FDP) when using $s_{j}$ as a cutoff. In our procedure, we use an estimated null distribution, and we adjust the estimated FDP by our estimated proportion of null hypothesis, ${\overset{̂}{\pi}}_{0}$. We also note that the number of tests $M$ is random in the DRP clustering problem, so for formal FDR control one might assume the cluster scores $s_{j}$ are independent conditional on $M.$

5. **Breakpoint intervals**

Discordant read clusters alone do not give precise breakpoint locations, so we provide an estimated breakpoint location along with a confidence interval. A natural estimate for the breakpoint position uses the locations of the $3^{'}$-most aligned bases among all reads in the cluster. For example, a deletion cluster with $3^{'}$ positions $\{a_{i}\}$ and $\{b_{i}\}$ yields breakpoint estimates $\max_{i}a_{i}$ and $\min_{i}b_{i}$. The true distribution of $3^{'}$ positions around each breakpoint will depend on the insert size distribution. Here, we estimate using a uniform distribution on an unknown but fixed genomic interval $[\underset{̲}{\theta},\overline{\theta})$ and use a 95% confidence interval for the breakpoint position, which we derive below.

Let $X_{1},\ldots,X_{n}$ be drawn i.i.d. from a Uniform distribution on $[a,b)\subset\mathbb{R}$. We construct a confidence interval for $a$ having the form

$$[X_{(1)}-c(X_{(n)}-X_{(1)}),X_{(1)}],$$

where $c>0$ depends on $n$ and the confidence level $1-\alpha$. To solve for $c$ we reduce the problem to one based on Uniform$(0,1)$ order statistics $U_{(1)},\ldots,U_{(n)}$. Defining

$$Z_{j}=\frac{\sum_{i=1}^{j} E_{i}}{\sum_{i=1}^{n+1} E_{i}},$$

where $E_{i}$ are independent Exponential$(1)$ random variables, it is known that $(U_{(1)},\ldots,U_{(n)})$ and $(Z_{1},\ldots,Z_{n})$ have the same joint distribution (DasGupta 2011). Now we have

$$\begin{matrix} 1-\alpha= & P\left( a\geq X_{(1)}-c(X_{(n)}-X_{(1)}) \right) \\ = & P\left( \frac{a-X_{(1)}}{X_{(n)}-X_{(1)}}\geq c \right) \\ = & P\left( \frac{-U_{(1)}}{U_{(n)}-U_{(1)}}\geq c \right) \\ = & P\left( \frac{-E_{1}}{\sum_{i=2}^{n} E_{i}}\geq c \right). \end{matrix}$$

The final term concerns Exp(1) divided by an independent Gamma($n-1$, 1), which by definition follows a scaled F-distribution: $\frac{2}{2n-2}F_{2,2n-2}$. Thus, $c$ can be obtained using the appropriate quantile function. A confidence interval for the upper endpoint $b$ is given by the same argument.

6. **Breakpoint merging**

After the above steps, we have breakpoint locations derived from soft-clipped reads, split reads, and discordant read pairs. We merge all overlapping breakpoint intervals and exact breakpoints (represented as intervals of 1 bp) and keep merged intervals with a supporting DRP cluster or at least 2 supporting split/clipped reads. Given merged breakpoint intervals $\{I_{j}=[a_{j},b_{j}){\}}_{j=1}^{n_{\mathrm{bp}}}$, along with $I_{0}=[0,1)$ and $I_{n_{\mathrm{bp}}+1}=[L,L+1)$, where $L$ is the length of the chromosome, we define blocks $B_{j}=[b_{j-1}-1,a_{j})$ for $j=1,\ldots,n_{\mathrm{bp}}+1$. (Note that if there is breakpoint uncertainty, i.e., if $b_{j}-a_{j}>1$ for some $j$, then there will be gaps between some adjacent blocks.) These blocks are the genomic segments that will be rearranged during SV detection.

***Construction of Adjacency Graphs***

The adjacency graph is an undirected multigraph used to represent adjacencies between genomic segments in the sample genome. Each segment $B_{j}$ in the reference genome is given two nodes, $B_{j}^{\mathrm{in}}$ and $B_{j}^{\mathrm{out}}$, corresponding to the start and end of $B_{j}$ relative to the reference coordinates; insertion blocks $S_{i}$ are treated likewise. We add “block edges” between each block’s pair of nodes while all other edges are defined as “adjacency edges,” as they encode the information that two blocks may be adjacent in the sample genome. Note our construction is technically a multigraph because there may be both a block edge and adjacency edge between $B_{j}^{\mathrm{in}}$ and $B_{j}^{\mathrm{out}}$ as in the case of a tandem duplication $B_{j}B_{j}$. The block edge merely represents the genomic segment $B_{j}$, while adjacency edge represents the novel adjacency introduced by $B_{j}B_{j}$.

1. **Construction**

Adjacency edges $B_{j}^{\mathrm{out}}-B_{j+1}^{\mathrm{in}}$, which are implied by the reference sequence, are added automatically, as are edges connecting candidate insertions $S_{i}$ to their flanking blocks. Other adjacency edges must be supported by the read alignments. For example, a read pair aligned to the forward strand of $B_{i}$ and to the reverse strand of $B_{k}$ is considered to support the edge $B_{j}^{\mathrm{out}}-B_{k}^{\mathrm{in}}$ if the implied insert size under that adjacency falls within the expected range (middle 95% of the insert distribution). An exception is made for adjacencies $B_{i}^{\mathrm{out}}-B_{i}^{\mathrm{in}}$, i.e., those supporting a tandem duplication of a single block. In this case, we additionally require that insert size (possibly negative) implied under no tandem duplication falls outside the acceptable 95%. Split reads whose splits occur at candidate breakpoints automatically add support to the implied novel adjacencies. Finally, note that we do not include any adjacency edges across gaps in the reference assembly. The final adjacency graph is formed by removing edges with only 1 supporting read, as well as edges spanning more than 2 Mb. This distance is longer than, for example, the largest deletion in the HuRef Gold Set (Mu et al. 2015).

2. **Partition and traversal**

Paths through the adjacency graph will correspond to candidate SVs. To avoid a combinatorial explosion in the number of paths, we divide the adjacency graph into a set of subgraphs (Supplementary Figure S11). We call a reference block $B_{j}$ “spanned” by the adjacency graph if there is some edge connecting $B_{i}$ to $B_{k}$, where $i<j<k$. We define a “back edge” from $B_{j}^{\mathrm{out}}$ as any edge connecting to a block $B_{i}$ with $i\leq j$; similarly, a “back edge” from $B_{j}^{\mathrm{in}}$ connects $B_{j}^{\mathrm{in}}$ to some block $B_{k}$ with $k\geq j$. Intuitively, if one first crosses a genomic segment using a block edge, then a “back edge” is any edge that causes the path to “change direction”. We call $j$ a cut point in case $B_{j}$ is not spanned and there are no back edges connected to $B_{j}^{\mathrm{in}}$ or $B_{j}^{\mathrm{out}}$. Additionally, the first and last reference blocks, as well as blocks adjacent to assembly gaps, are considered cut points. Thus, we have cut points $1=j_{1}<j_{2}<\cdots<j_{P}=J$ within the graph. For each two consecutive cut points $j_{m}$ and $j_{m+1}$, we will consider the subgraph formed by blocks $B_{j_{m}},\ldots,B_{j_{m+1}}$ as well as any insertion blocks contained within that genomic region. Subgraphs consisting of only a pair of reference blocks are discarded. Finally, each subgraph is extended on either side to ensure that some minimal amount of sequence flanks any SV, and subgraphs that overlap as a result are merged.

Given two nodes $s$ and $e$, a valid traversal through the graph is a path from $s$ to $e$ that alternates between block edges and adjacency edges, beginning and ending with block edges. Any valid traversal is equivalent to some sequence of oriented blocks, i.e., a genomic rearrangement. For tractable computation, we limit the number of times (by default, 2) each node may be exited using any back edges. Note that we have implicitly limited the number of cycles in a valid traversal. Limiting only the multiplicity of cycles in the path may rapidly expand the number of paths around large tandem duplications.

***Likelihood model***

Suppose a subgraph, or region $R$, consists of reference blocks $B_{i},B_{i+1},\ldots,B_{j}$. Let $\Theta$ denote the set of valid traversals from $B_{i}^{\mathrm{in}}$ or $B_{j}^{\mathrm{out}}$, so that each $\theta\in\Theta$ represents a possible rearrangement of $R$. The set of genomic coordinates within $R$ under rearrangement $\theta$ is called $\mathcal{R}_{\theta}$. This set is $[0, L_{\theta})$, where $L_{\theta}$ is the length of the rearranged version of $R$. Now, we aim to maximize the likelihood of the observed paired-end alignments, $P(\mathrm{data};\theta_{1},\theta_{2})$, jointly over the sample’s diploid genotype $(\theta_{1},\theta_{2})\in\Theta^{2}$.

1. **Fragment and alignment model**

Sequenced DNA fragments are sampled from $\mathcal{R}_{\theta}$ with frequencies proportional to the insert size density. If $x,y\in\mathcal{R}_{\theta}$ and $x<y$, fragment $[x,y)$ is sampled with probability

$$P(x,y;\theta)=\frac{f(y-x)}{\sum_{u,v\in\mathcal{R}_{\theta},u<v} f(u-v)}.$$

After the paired-ends of fragment $[x,y)$ are sequenced, we observe a paired-end alignment to the reference genome, $\underset{̲}{a}=(a_{1},a_{2})$. Specifically, the alignment $a_{j}$ is represented as $a_{j}=(s,o,m)$. The variable $s$ is the sequence of oriented reference blocks corresponding to the alignment. For example, a read from the forward strand of $B_{1}$ has $s=(B_{1})$, whereas a read from the reverse strand has $s=(B_{1}^{'})$; a split alignment supporting the deletion of $B_{2}$ may have $s=(B_{1},B_{3})$. We use $o$ to denote the read’s start position, or offset, within the first block of $s$ where $s$ and $o$ are directly observed (Supplementary Figure S12a). We do not explicitly model soft-clipping: clipped bases on the $3^{'}$ end are ignored, and reads with clipped $5^{'}$ ends have their offsets $o$ adjusted so the fragment length is calculated correctly.

The variable $m$ indicates the alignment outcome of the sampled read. Ordinary paired-end alignments $(a_{1},a_{2})$ to the region $R$ have $m_{1}=m_{2}=1$. A “hanging pair” in which only read 1 is aligned is indicated by $m_{1}=1,m_{2}\neq1$ (Supplementary Figure 13a). In this case we then set $m_{2}=0$ if read 2 is unmapped and $m_{2}=-1$ if read 2 is mapped outside of $R$. Pairs in which neither read is mapped to $R$ (i.e., $m_{1}\neq1$ and $m_{2}\neq1$) are assumed unobserved. Thus, we perform inference conditional on the complementary event, $m_{1}=1\vee m_{2}=1$.

Here, the frequency of hanging and unmapped reads depends on the unobserved variable $\underset{̲}{I}=(I_{1},I_{2})$, where $I_{j}$ is 1 if read $j$ is derived from a novel insertion and 0 if it derives from reference sequence. For the case where both reads derive from reference sequence, we estimate the distribution $P(m_{1},m_{2}|I_{1}=I_{2}=0)$ using the observed frequencies of mapped and hanging pairs across all reads. This is a conservative overestimate since we include insertion-derived reads. To handle reads within insertions, we assume the probability of successfully mapping a reference-derived read does not depend on the location of its mate:

$$P(m_{1}=1|I_{1}=0,I_{2}=1)=P(m_{1}=1|I_{1}=0,I_{2}=0).$$

Conditional on the reference-derived read aligning successfully ($m_{1}=1$ above), we assume the insertion-derived read is either unmapped or mapped distantly ($m_{2}=0,-1$) with equal probability. If the one reference-derived read is unmapped, or if both reads derive from inserted sequence, then we assume neither $m_{1},m_{2}$ is equal to 1, and the pair is not observed.

2. **Likelihood evaluation**

Given a paired end alignment $\underset{̲}{a}=(a_{1},a_{2})$ and a candidate haplotype $\theta$, we consider the set of fragments from $\mathcal{R}_{\theta}$ that have block sequences and offsets identical to those of $(a_{1},a_{2})$. We call this set of “compatible” fragments $C_{\theta}(\underset{̲}{a})$. Each member of $C_{\theta}(\underset{̲}{a})$ is an interval $[x,y)\subseteq\mathcal{R}_{\theta}$ and a variable $\xi\in\{1,2\}$, which indicates which one of $a_{1}$ or $a_{2}$ corresponds to the left end of the fragment (in the coordinates of the sample genome). For example, suppose read 1 matches the forward strand of $B_{1}$ and read 2 matches the reverse strand of $B_{3}$, then there is 1 compatible fragment under $\theta=B_{1}B_{3}$, none under $\theta=B_{1}B_{2}$, and 2 under $\theta=B_{1}B_{2}B_{3}B_{3}$. If only one read is mapped, then $C_{\theta}(\underset{̲}{a})$ consists of all fragments matching the mapped end (Supplementary Figures S12b and S13b).

We assume that $\underset{̲}{a}$ is observed conditional on the mapped end(s) falling within the region $R$. Summing over the compatible fragments and incorporating the hanging read model, we have the likelihood

$$\begin{matrix} P_{\theta}(\underset{̲}{a}|mapped ends in \mathcal{R}_{\theta})= & \frac{\sum_{([x,y),\xi)\in C_{\theta}(\underset{̲}{a})} f(y-x)P\left( \underset{̲}{m}|\underset{̲}{I}=\eta_{\theta}([x,y),\xi) \right)}{\sum_{x,y\in\mathcal{R}_{\theta}:x<y} \sum_{\xi\in\{1,2\}} f(y-x)P\left( m_{1}=1\vee m_{2}=1|\underset{̲}{I}=\eta_{\theta}([x,y),\xi) \right)} \\ \equiv& \frac{p_{\theta}(\underset{̲}{a})}{G_{\theta}}, \end{matrix}$$

where $\eta_{\theta}$ indicates whether the reads derive from novel insertion sequence.

If the rearrangement $\theta$ contains no novel insertions, then the normalizing constant $G_{\theta}$ merely requires computing a double sum $\sum_{x=0}^{L} \sum_{y=x+1}^{L} f(y-x)$, where $L$ is the length of the haplotype. Adjusting for $N$ novel insertions requires an additional computation of $O(N^{2})$ similar double sums. Each such term is computed in $O(1)$ time using pre-computed values of $F(t)=\sum_{x=0}^{t} f(x)$ and $\overline{F}(t)=\sum_{x=0}^{t} F(x)$.

The likelihood of the data, assuming fragments are sampled independently, is then

$$P_{\theta}(\mathrm{data})=\frac{1}{G_{\theta}^{n}}\prod_{i=1}^{n} p_{\theta}({\underset{̲}{a}}_{i}).$$

Note that $p_{\theta}({\underset{̲}{a}}_{i})=0$ if ${\underset{̲}{a}}_{i}$ has no compatible fragments under $\theta$. This may occur even if $\theta$ is correct, for example if there is a spurious split alignment or an error in mapping orientation. Thus, we use the following “robust” likelihood,

$${\overset{̃}{P}}_{\theta}(\mathrm{data})=\frac{1}{G_{\theta}^{n}}\prod_{i=1}^{n} (\pi+(1-\pi)p_{\theta}({\underset{̲}{a}}_{i})).$$

This modified likelihood gives a small probability to all possible mappings, eliminating the effect of extreme outliers. Here, we choose the robustness parameter ⇡ using the insert size distribution:

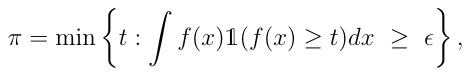

where **1** is the indicator function. For a symmetric unimodal density $f$ with cdf $F$, we have $\pi=f(F^{-1}\left( \frac{\epsilon}{2} \right))$. In practice, we evaluate the expression above with $\epsilon=2\Phi(-4)$, so that extreme outliers in our model are about as surprising an observation of ±4 from the normal distribution.

3. **Computational considerations**

Since we are working with diploid genomes, in which region $\mathcal{R}$ exists as a pair of (possibly identical) haplotypes $(\theta_{1},\theta_{2})\in\Theta$. We do not know from which haplotype each read pair is sampled, so we must use the following mixture likelihood:

$${\overset{̃}{P}}_{\theta_{1},\theta_{2}}(\mathrm{data})=\frac{1}{\left( G_{\theta_{1}}+G_{\theta_{2}} \right)^{n}}\prod_{i=1}^{n} (2\pi+(1-\pi)(p_{\theta_{1}}({\underset{̲}{a}}_{i})+p_{\theta_{2}}({\underset{̲}{a}}_{i})).$$

This likelihood function is derived under the assumption that each haplotype is sampled with probability proportional to its length, which is reasonable if we are sampling a single genome with paired chromosomes. If sequencing data were collected from a pool of individuals, one could adjust the above likelihood so that an unknown pro- portion ⇢ of chromosomes have haplotype $\theta_{1}$, while the rest have haplotype $\theta_{2}$.

To summarize, we use the following procedure in evaluating and maximizing the likelihood:

Specifically, we use the following procedure:

i. Enumerate rearrangements $\theta_{1}\in\Theta$ based on traversals through the current subgraph $R$. If the size of $\Theta$is greater than 1000, skip this subgraph.
ii. For each $\theta$, evaluate the likelihood under $(\theta_{\mathrm{ref}},\theta)$ and $\left( \theta,\theta\right)$, where $\theta_{\mathrm{ref}}\in\Theta$ is the reference haplotype. Rank each $\theta_{\mathrm{ref}}$including $\theta_{\mathrm{ref}}$ according to the higher of the two likelihoods and let $\bar{\Theta}\subseteq\Theta$ contain the top 50 $\theta$.
iii. Maximize the diploid likelihood ${\overset{̃}{P}}_{\theta_{1},\theta_{2}}$ over $(\theta_{1},\theta_{2})\in\bar{\Theta}^{2}$ to produce the final SV call.

### ***Implementation details for calling SVs***

Recall that there may be gaps between some blocks due to breakpoint uncertainty. It is possible (though rare in practice) that an alignment falls entirely into one of these gaps; these reads are ignored. Also, when rearranging blocks, we take the median of any breakpoint gap to be the best guess for the breakpoint location. For example, a deletion of $B$ from $ABC$ will leave behind half of any gaps between $A$ and $B$, $B$ and $C$.

Our algorithm only uses aligned reads with a MAPQ score of at least 20, except when determining candidate breakpoints from soft-clipped reads (at least 10). The unmapped or “distantly mapped” ends of hanging reads are not subject to a MAPQ constraint. Split reads with breakpoint positions that do not agree with our merged, filtered breakpoints are ignored.

***Affected sequence calculation***

A simple SV is defined by a single position and length in the reference genome. Complex SVs, however, may span large regions but make only small modifications to the sequence. In our analysis of complex SV calls, we keep the usual requirement that SVs must affect at least 50 bp of sequence. We propose a simple method that defines which reference segments are “affected” by an SV.

Suppose the reference region (separated into segments by the breakpoints) is given by $B_{1}^{0}B_{2}^{0}\ldots B_{n}^{0}$, and the rearranged version is $B_{r_{1}}^{o_{1}}B_{r_{2}}^{o_{2}}\ldots B_{r_{m}}^{o_{m}}$, where $o_{i}$ is 1 if the segment is in reverse orientation. We find a pairwise alignment between these two strings of genomic segments (not the nucleotide sequences), treating blocks $B_{i}^{o_{i}}$ and $B_{j}^{o_{j}}$ as equal if $i=j$ and $o_{i}=o_{j}$. Alignments were computed using a very large score (1000) for matches and equally small penalties (-1) for mismatches and gap extensions. After finding an optimal alignment, blocks contained in either sequence that are unaligned or mismatched are considered “affected.” The affected length is then computed as the sum of affected reference block lengths. We used the Biostrings::pairwiseAlignment function in R (<https://bioconductor.org/packages/Biostrings>), which returns only a single optimal alignment. To partially account for non-unique optimal alignments, we computed a second alignment by switching the subject and query sequences and adding any additional affected blocks. For additional discussion of background, motivation, and implementation of the ARC-SV probabilistic model, see J. Arthur Doctoral Thesis (Arthur 2018)

***Computational validation of SVs***

We validate SV predictions by direct sequence comparison to ground truths i.e. high-quality diploid genome assemblies (Liao et al. 2022). The SV is considered validated if the predicted sequence matches the truth. We construct a predicted sequence for each cxSV and SV call by rearranging the reference genome accordingly for each SV called and included up to 1kb of flanking sequence on each side (software default). This altered sequence is annotated with the locations and sizes of any deletions as well as the boundaries of the rearranged reference blocks. We say block $B_{k}$ is deleted if $B_{k}$ is absent in the rearrangement and the subsequence $B_{k-1}B_{k+1}$ is present in either orientation.

Before alignment, prediction sequences are modified to such that only 150 bp of flanking sequences remain. This is so that flanking sequences serve only the role of anchor and have minimal effect on the overall alignment score of the prediction sequence. Following the validation paradigm developed for detecting cxSVs in long-reads (Sedlazeck et al. 2018), a SV prediction sequences is validated if its sequence prediction of rearranged genome segments can be continuously aligned in its entirety from beginning to end (including both flanking anchor sequences) i.e. 100% “query coverage”, without any soft or hard-clipping, with maximum mapping quality score (mapq=60), with >90% BLAST identity (Altschul et al. 1990) to the ground truth and additionally with an absolute maximum alignment gap of no larger than 49 bp to allow possible INDEL variants and breakpoint interval uncertainty in the cases of mostly DRP support for the prediction sequence. Alignments were made using minimap2 v2.18 using default parameters (Li 2018).

**Machine learning model for obtaining high-confidence calls**

Next, to identify high-confidence SV calls, we optimized machine learning models (random forest, XGBoost, lightGBM) for each SV type (cxSVs, deletions, tandem duplications, and inversions) implemented using the scikit-learn package version 1.1.2 (<https://scikit-learn.org>) for Python (version 3.8.2). Our model predicts validation of SV that are given 48 features transformed from the output file of ARC-SV probabilistic model (Supplementary Table S4) where categorical information in “rearrangement block configuration” and “reference block configuration” are pre-processed as matrices (Supplementary Figure S14). Matrices are created using a mapping of vectorized reference blocks and rearrangement blocks. These matrices are composed of binary numbers (0 or 1). Since these matrices are sparse, we decompose them using Principal Component Analysis (PCA) and reduced to $\mathbb{R}^{N}\times\mathbb{R}^{2}$ dimensional matrices (Supplementary Figure S14).

For training and test sets, we used 42 diploid genome assemblies from the Year 1 Release of the Human Pangenome Reference Consortium (Liao et al. 2022) where 31 (74%) genomes were used for training and 11 (26%) were used for testing the machine learning model (Supplementary Table S3,S4). The split of training and test sets were stratified by genome ancestry. SVs calls from each genome based on ~30x-coverage paired-end WGS were validated against the diploid genome assemblies (see Computational Validation of SVs section) and labeled based on validation outcome. Specifically, for each SV class (deletion; tandem duplication; inversion; complex), we have features $x_{i}$ and a label $y_{i}\in\left\{ 0,1 \right\}$ indicating whether the SV is a “non-validated” (False) or “validated” (True) detection.

We applied three machine learning approaches (random forest, extreme gradient boost, and lightGBM) to identify validated cxSV and simple SV calls. For cxSVs, the optimal machine learning model based on maximum precision score is a XGBoost model (n_estimators=160, max_depth=143, gamma=0, random_seed=559) (Supplementary Table S5) achieving 95.7% precision and 91.4% recall) whereas models for inversions (random forest: bootstrap=True, max_depth=215, max_features='auto', min_samples_leaf=1, min_samples_split=2), duplications (XGBoost: base_score=0.5, gamma=0, learning_rate=0.3, max_depth=14, min_samples_split=14, n_estimators=195), and deletions (XGBoost: base_score=0.5, gamma=0, learning_rate=0.3, max_depth=84, min_samples_split=14, n_estimators=360) were chosen (precision/recall: 96.3%/93.1%, 94.7%/90.6%, and 97.2%/96.8% respectively) (Supplementary Table S5).
